# Plumage balances camouflage and thermoregulation in Horned Larks (*Eremophila alpestris*)

**DOI:** 10.1101/2021.07.15.452373

**Authors:** Nicholas A. Mason, Eric A. Riddell, Felisha Romero, Carla Cicero, Rauri C.K. Bowie

## Abstract

Animal coloration serves many biological functions and must therefore balance potentially competing selective pressures. For example, many animals have camouflage, in which coloration matches the visual background against which predators scan for prey. However, different colors reflect different amounts of solar radiation and may therefore have thermoregulatory implications as well. In this study, we examined geographic variation in dorsal patterning, color, and solar reflectance among Horned Larks (*Eremophila alpestris*) of the western United States. We found associations between dorsal plumage brightness, hue, and patterning relative to the soil conditions where specimens were collected. Specifically, brighter dorsal plumage corresponded to brighter soil, while redder, more saturated hues in dorsal plumage corresponded to redder soils. Furthermore, backs with more high-contrast patterning were more common among females and also associated with soil that had coarser soil fragments, suggesting that lark plumage has been selected to optimize background matching in different environments. We also found that larks exhibited higher solar reflectance in hotter and more arid environments, which lowers the water requirements for homeothermy. Taken together, these findings suggest that natural selection has balanced camouflage and thermoregulation in Horned Larks across a wide variety of soil types and abiotic conditions.

## Introduction

Animal colors and patterns constitute complex phenotypes that are shaped by a wide array of biotic and abiotic processes (Burtt 1981; Vo et al. 2011; Cuthill et al. 2017). For example, some species have bright colors involved in sexual selection via mate choice (Andersson and Simmons 2006; Shultz and Burns 2017), whereas others have cryptic colors and patterns driven by natural selection to avoid visual detection by predators (Cott 1944; Endler 1978). Furthermore, animal colors have implications for maintaining homeostasis in different environments due to differences in the reflectance of light wavelengths from solar radiation (Walsberg 1983; Wolf and Walsberg 2000). Thus, animal coloration and patterning must balance multiple selective pressures that may conflict or act synergistically to produce multifunctional phenotypes that vary among populations and species (Caro 2017). Among the wide array of biological processes affecting animal colors, camouflage and thermoregulation are thought to have a particularly strong influence on coloration because of their direct impact on survival.

Camouflage includes a suite of physical and behavioral attributes that deter visual detection by predators and is prevalent among various animal lineages, including arthropods (Farkas et al. 2013; Stevens et al. 2014) and vertebrates (Rosenblum et al. 2009; Isaac and Gregory 2013; Boratyński et al. 2017). Also known as background matching, camouflage favors phenotypes that resemble a random sample of brightness, hue, and patterning of the visual background against which predators actively scan for prey (Endler 1978; Merilaita et al. 1999; Michalis et al. 2017). Thermoregulation is also closely tied to coloration (Walsberg 1983) and sometimes favors phenotypes that either conflict with background matching (Smith et al. 2016) or that simultaneously enable thermoregulation and camouflage (Wacker et al. 2016). Many species conform to ‘Gloger’s Rule,’ a highly prevalent ecogeographic pattern that ascribes lighter colors to more xeric environments and darker colors to more mesic environments (Gloger 1833; Delhey 2019; Marcondes et al. 2020). This widespread trend is thought to be the product of multiple selective pressures, including camouflage and thermoregulation (Burtt 2004; Delhey et al. 2019). Plumage reflects and absorbs light that includes wavelengths within the visual range of birds (UV-VIS: 300–700 nm) as well as near-infrared wavelengths (NIR: 700–2,600 nm), both of which serve important roles in light and heat absorptance (Stuart-Fox et al. 2017). Within the visual spectrum, darker feathers tend to absorb more light and heat than lighter feathers (Porter and Gates 1969), but the physical properties of feathers and the ability of incident light to travel through or become captured by feather microstructures also play important roles (Walsberg 1988; Wolf and Walsberg 2000). Because NIR wavelengths are not perceived by predators, reflectance at those wavelengths is not related to camouflage but may still play an important thermoregulatory role (Medina et al. 2018).

Despite the prevalence of camouflage among animals, the majority of studies to date on background matching have focused on a small number of systems such as peppered moths (Van’t Hof et al. 2011; Cook and Saccheri 2013), pocket mice (Nachman et al. 2003; Linnen et al. 2009), White Sands lizards (Rosenblum et al. 2010; Laurent et al. 2016), and a few ground-nesting birds (Troscianko et al. 2016; Stevens et al. 2017). Most of these systems involve discrete phenotypic variants that occupy visually distinct environments. In comparison, continuous variation in background matching across environmental gradients has received far less attention (Stevens and Merilaita 2009; Caro et al. 2016). Furthermore, studies that simultaneously examine camouflage and thermoregulation remain scarce, especially among endotherms, such as birds. Finally, the role of NIR reflectance in potentially mediating tradeoffs between camouflage and thermoregulation remain largely unexplored (Stuart-Fox et al. 2017; Medina et al. 2018).

To address these knowledge gaps, we examined associations between plumage reflectance and patterning, soil color and composition, and thermoregulatory models among geographically variable populations in a widespread songbird, the Horned Lark (*Eremophila alpestris*). Horned Larks occupy a wide variety of open mesic and arid habitats, including deserts, fallow agricultural land, tundra, and grasslands (Beason 1995; Mason et al. 2020). Horned larks build nests and glean seeds and insects on the ground (Wiens and Rotenberry 1979; de Zwaan and Martin 2018). Due to their preference for habitats with sparse vegetation, larks are thought to rely on substrate matching to avoid avian predators (Donald et al. 2017). Although camouflage in Horned Larks has been discussed anecdotally (Zink and Remsen 1986; Mason and Unitt 2018), associations between phenotypic and environmental variation have not yet been tested rigorously. Larks also exhibit various physiological adaptations to aridity gradients (Tieleman et al. 2003*b*, 2003*a*), making them an excellent system to study interactions between camouflage and thermoregulation. If larks exhibit background matching, we predict that soil conditions will be associated with variation in plumage brightness, color, and patterning. Furthermore, if plumage also plays a thermoregulatory role, we predict that rates of evaporative water loss and solar reflectance will be associated with variation in aridity and temperature.

To test these predictions, we combined digital photography, color and plumage analyses, full-spectrum (UV, Visual, NIR) spectroradiometry of museum specimens, remote sensing data, and simulation-based thermoregulatory models of heat flux to examine phenotype-environment associations between plumage coloration and patterning, soil conditions, and climate. This approach allows us to disentangle the effects of camouflage, thermoregulation, and sexual dimorphism in driving the evolution and ecology of lark coloration. More broadly, it illustrates how we can understand limits on the adaptive potential of certain traits such as coloration. For example, warming climates might select for more reflective features, but at a cost to background matching. Likewise, habitat alterations might select for darker feathers that impose greater physiological stress under warming climates. Thus, one selective pressure might have detrimental effects to another in the evolution of phenotypes. Only by integrating both background matching and thermoregulatory performance can we understand evolutionary responses to these different and potentially competing selective pressures.

## Methods

### Digital Photography and Image Analysis

We photographed the dorsal side of 270 Horned Lark specimens from the Museum of Vertebrate Zoology (MVZ) at the University of California, Berkeley (Supplementary Table S1), using a Nikon D7000 camera modified for full-quartz calibration (Advanced Camera Services, Watton, Norfolk, England). We measured up to 10 males and 10 females of 17 different subspecies (Figure 1) in the western United States, preferentially selecting specimens from breeding months (May–August) and with undamaged plumage. We used a Novoflex Noflexar 30mm f/3.5 lens, which does not filter out ultraviolet wavelengths and is therefore suitable for measuring plumage reflectance under an avian visual model. We took two RAW images of each specimen at ISO200: one image used a Baader Venus-U filter, which captures wavelengths between ∼320–380 nm, and a second image used a Baader UV/IR cut filter, which captures wavelengths between ∼400–680 nm. Each image included a ruler at the height of the specimen’s dorsal plane with 5% and 80% reflectance standards (Labsphere, Hutton, NH, USA). We automatically aligned and linearized images using the Image Calibration Analysis Toolbox (Troscianko and Stevens 2015), which provides a set of plugins for ImageJ (Schneider et al. 2012), and manually drew polygons corresponding to the dorsal region of each specimen in ImageJ (Supplementary Figure S1).

**Figure 1:**
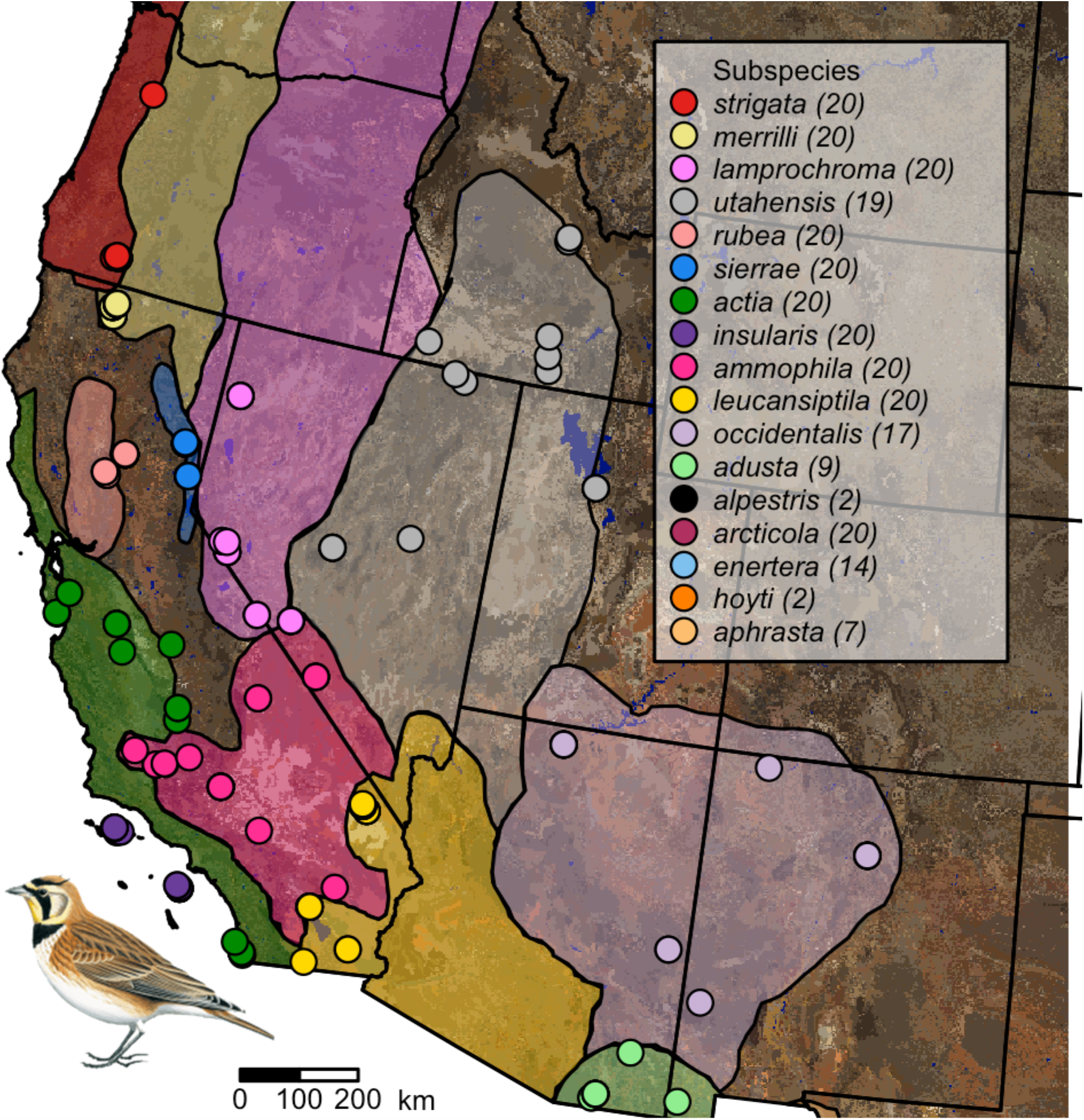
Sampling map showing localities of vouchered Horned Lark (*Eremophila alpestris*) specimens used in this study. Soil color is based on USDA soil surveys, and the approximate range of each subspecies is shown in a different color based on Behle (1942) and new museum records. Some dots may represent more than one individual sampled from the same locality. The number of samples per subspecies is given in parentheses next to the name of the subspecies in the legend in the upper right. Some subspecies (*alpestris, arcticola, enertera, hoyti, aphrasta*) are not shown because we focused on the geographic area with the most dense sampling.

After processing each image and delimiting the dorsal region of interest, we converted the channel readings for the UV and visual images to the cone-catch values of a blue tit visual model (Vorobyev and Osorio 1998). We then converted these cone-catch values into tetracolorspace measurements (Vorobyev et al. 1998; *sensu* Stoddard and Prum 2008) of hue, saturation, and chroma using the package pavo v2.2.0 (Maia et al. 2013, 2019) in the R programming environment (R Core Team 2020). We also measured achromatic brightness (i.e., total reflectance or luminosity across all wavelengths) and calculated an index of patterning via a series of Fast Fourier Transform (FFT) bandpass filters at 49 levels (beginning at 2 pixels and increasing exponentially by 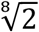 to 128). FFTs are widely applied in digital image analysis (Stoddard and Osorio 2019) and can be used to quantify animal patterns based on neurophysiological processing of spatial patterns (Godfrey et al. 1987; Stoddard and Stevens 2010; Troscianko et al. 2016; Mason and Bowie 2020). In this process, an image is converted into a set of sine waves, each with a different frequency and amplitude. The amplitude or power of each wave indicates how much patterning—or change between light and dark pixels—occurs at a specific spatial scale. Thus, images with high power across FFT bandwidths display more patterning (e.g. the dorsal spots on some larks), whereas images with lower power are more uniformly colored.

Finally, we performed a principal component analysis on the three focal plumage characters: brightness, achieved chroma, and back patterning (total power) to summarize plumage variation among the sampled larks. We found that the first principal component axis loaded positively with brightness and dorsal patterning, whereas the second principal component axis represented a tradeoff between brightness and back patterning (Supplementary Table S2).

### Measuring Full-spectrum Solar Reflectance

We measured solar reflectance of the lark specimens described above using recently published methods for estimating heat stress in birds (Riddell et al. 2019). For each specimen, we measured dorsal and ventral feather reflectance from 350 – 2500 nm using an ASD FieldSpec Pro spectroradiometer (ASD, Inc., 1625 S. Fordam Street, Suite 300, Longmont, CO 80503), and standardized each measurement relative to a Spectralon™ white standard before recording our measurements. We used a custom-built tungsten halogen light source to measure feather reflectance 2 cm from the feather surface, with a 45° angle between the light source and fiber optic cable. This light source was built using an AC-to-DC voltage converter to inhibit interference from an alternating source of electrical current. To standardize the angle and distance, we used a RPH-1 reflection probe holder (Ocean Optics, Inc., Largo, FL). We also used the reflection probe to standardize the surface area of each measurement to ensure that the measurements were not influenced by body size. The combination of numerous measurements per specimen (see below) and the diameter of the probe opening (6.35 mm) ensured that we captured the average solar reflectance of each side for each specimen.

We used ViewSpec software (ASD, Inc.) to measure solar reflectance, recording ten measurements (five dorsal and five ventral) for each of the 270 specimens. On both the dorsal and the ventral sides, we recorded one measurement from the crown or neck and four measurements spread across the breast or mantle for a total of 2,700 measurements. We then used a custom script in Python (v. 3.5) to average these values for each individual, and corrected the reflectance curves for solar radiation using the ASTM G-172 standard irradiance spectrum for dry air provided by SMARTs v. 2.9 (Gueymard 2001). We calculated the corrected value by multiplying the intensity of solar radiation by the empirical reflectance, integrating across all of the wavelengths, and dividing by the total intensity of solar radiation (Gates 1980).

### Remote Sensing Data

We compiled two different soil data sets to examine associations between lark dorsal plumage and soil conditions. First, we downloaded a soil color data set based on an extensive series of United States Department of Agriculture soil surveys of the contiguous United States, with values that had been converted from Munsell color charts to RGB color space (Beaudette et al. 2013). Using georeferenced localities of each lark specimen obtained from the MVZ database Arctos (arctos.database.museum), we extracted their respective soil values and performed a principal components analysis to assess soil color. The first principal component axis loaded strongly with all three channels corresponding to soil brightness. The second principal component axis loaded positively with the red channel, but negatively with blue and green (Supplementary Table S3), and therefore corresponded to soil redness. In this manner, we obtained soil color data associated with the site of collection of 224 of our lark specimens. This soil color dataset is limited to the continental United States, so we were unable to include 46 individuals from populations in Alaska, Canada, and Mexico in this part of our analysis.

We also downloaded harmonized soil property data for the top five centimeters of soil depth at a 30 arc-second resolution from the WISE30Sec database (Batjes 2016). WISE30Sec data has been used widely to study soil biogeochemistry for quantifying global carbon stocks (Sanderman et al. 2017). Although it has not been applied broadly to organismal biology, this dataset provides ecologically relevant information on clay abundance, proportion of coarse fragments, and other soil properties relevant to terrestrial organisms such as Horned Larks. To generate an index of soil surface granularity, we extracted the volume percentage of coarse fragments (> 2 mm) and the mass percentages of sand, silt, and clay. We then performed a principal component analysis and found that the first principal component axis loaded positively with coarse fragments and sand, and negatively with silt and clay (Supplementary Table S4).

To examine associations between climate and plumage, we also downloaded all 19 WorldClim bioclimatic variables (worldclim.org; Hijmans et al. 2005) at a resolution of 30 arc-seconds to examine associations with dorsal plumage. We conducted a principal component analysis in which the first principal component axis loaded positively with seasonality, the second principal component axis loaded positively with aridity, and the third principal component axis loaded positively with temperature (Supplementary Table S4).

### Statistical Analyses of Phenotype-Environment Associations

We performed a series of statistical analyses to examine associations between dorsal plumage and the environment. First, we summarized phenotypic variation among subspecies by plotting the first two PCA axes of plumage variation and noted clustering by subspecies and sex. We then compared mean values by first performing an analysis of variance on the output of linear models, with subspecies as the grouping variable and with males and females separately. We subsequently used a Tukey’s multiple comparison post-hoc test (Steel et al. 1997) with the HSD test function from the agricolae package v1.3-3 (de Mendiburu 2020) in R (v4.0.1; R Core Team 2020) to assign subspecies to groups within each sex based on their mean values. We then constructed linear models (LMs) to quantify background matching by examining associations between plumage and soil variables with sex included as a main effect. Specifically, we constructed the following LMs: (1) plumage brightness as the response variable with soil brightness (soil color PC1) and sex as main effects; (2) plumage redness as the response variable with soil redness (soil color PC2) and sex as main effects; and (3) plumage patterning as the response variable with soil granularity (soil composition PC1) and sex as main effects. We also calculated Pearson’s product-moment correlation coefficients between plumage brightness and soil brightness (soil color PC1), plumage chroma and soil redness (soil color PC2), and plumage patterning (plumage patterning PC1) and soil granularity (soil granularity PC1) using the cor.test() function in R (v4.0.1; R Core Team 2020).

Finally, we generated additional LMs to simultaneously estimate the influence of soil and bioclimatic conditions on variation in plumage reflectance. Specifically, we constructed a LM with dorsal brightness (UV-VIS) as a response variable and soil color, seasonality, aridity, temperature, and sex as main effects using the glm() function in R (R Core Team 2020). We also generated a LM with dorsal solar reflectance (UV-VIS-IR) as a response variable and soil color, seasonality, aridity, temperature, and sex as main effects with the same function and settings.

### Heat flux simulations

We incorporated the empirical measurements of solar-corrected feather reflectance into a heat flux model to estimate the thermoregulatory differences among subspecies. This model simulates heat balance using the morphological characteristics of the bird in a complex radiative and thermal environment. The simulation output produces estimates of net sensible heat flux (Q), which was calculated using:

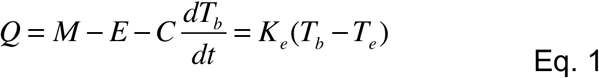

where *M* is the heat generated through metabolic processes, *E* is the heat lost via evaporative processes, *T*_*b*_ is body temperature, *K*_*e*_ is the effective conductance, and *T*_*e*_ is the operative temperature. The net sensible heat flux equation estimates the heat flux required to maintain a stable body temperature given the morphology of the bird and its interaction with the environment.

The heat flux simulation uses biophysical principles to estimate heat flux between the birds and their environment. We used environmental data generated by *NicheMapR* (v1.1.3; Kearney and Porter 2020) to estimate the thermal microclimate for larks. First, we obtained monthly minimum and maximum air temperatures from the *Worldclim* global climate database for the sites at which specimens had been collected. We then corrected these temperatures using *NicheMapR* to reflect conditions relevant to larks, using a reference height of 5 cm above the ground because larks spend most of their time near the ground. We simulated heat flux assuming two types of soil environments: a xeric, desert-like environment (sand, soil reflectance = 0.35) and a more mesic environment (loam, soil reflectance = 0.15; Campbell and Norman 1998). These two environments represent the extremes of the thermal environment that larks inhabit in our study area. Assuming a more reflective soil (i.e., sand) in our simulations did not qualitatively alter our conclusions. We then used these environments to understand the thermal consequences of variation in dorsal and ventral solar-corrected feather reflectance.

We incorporated morphological phenotypes that directly influence reflected solar radiation in several ways. Our goal was to isolate the importance of dorsal reflectance for thermoregulation. Thus, we assumed that each phenotype was equivalent among subspecies, with the exception of feather reflectance. In general, variation in body mass among subspecies of western Horned Larks is low (males = 29.7 g ± 2.4, females = 28.5 ± 3.5 g; Behle 1942), suggesting that the differences in mass are unlikely to substantially influence thermoregulatory differences among subspecies. Estimates of morphological phenotypes were taken from Riddell et al. (2019) using three specimens that represented the average mass of a lark in our simulations, but here we briefly describe the methods. We estimated plumage depth (sensu Kearney et al. 2016) by measuring the vertical distance from the skin to the outer surface of the feathers using a Fisherbrand™ 150 mm ruler at 10 locations that spanned the dorsal and ventral side of each specimen. We also measured the average length of contour feathers across six feathers per specimen spanning the dorsum and ventrum. To characterize the approximate shape of larks, we used measurements of the height, length, and width of the lark specimens. We measured length from crown to the vent, width from shoulder to shoulder, and height from the back of the dorsal side to the breast at the shoulder (Kearney et al. 2019). We then used these values to estimate the rough dimensions of a lark, assuming a spheroid shape (Porter and Kearney 2009). The dimensions of birds in nature are dependent upon posture and are thus highly variable. By using the same dimensions for each subspecies, our analysis focuses on the thermoregulatory effects of reflectance and avoids possible noise due to specimen preparation and behavioral differences among subspecies. The mean body mass for Horned Larks (29.5 g) was determined by Riddell et al. (2019), which used the Vertnet data aggregator (vertnet.org). Briefly, body mass was based on collection points in western North America (*n* = 2,468). The protocol in Riddell *et al*. (2019) removed data greater or less than two standard deviations from the mean body mass to remove juvenile values erroneously labelled as adult and extreme outliers that were likely a mistake. Estimates of mass agree closely with previously published values for subspecies of larks (Behle 1942).

The simulation estimates heat flux by integrating morphological phenotypes with environmental biophysics and behavior. Estimating heat flux in endotherms is complicated by properties of the insulation layer. We addressed these issues by integrating a series of equations involved in a two-dimensional heat transfer model to estimate the flux from the dorsal and ventral components of a bird (Bakken 1981). This model calculates the total amount of heat absorbed or lost from the environment, and converts the amount of energy into the physiological response that would be necessary to maintain a stable body temperature (39°C in our simulations). These values represent the amount of heat that needs to be generated via metabolic heat production or lost via evaporative cooling to regulate body temperature. We incorporated sources of heat specifically including air temperature, direct solar radiation, diffuse solar radiation, reflected radiation from the ground and sky, and longwave radiation from the sky and ground. We also calculated standard operative temperature (*T*_*es*_) to equate simulated environments in the field to laboratory conditions as shown in Equation 2:

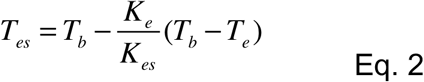

where *T*_*b*_ is body temperature, *K*_*e*_ is effective conductance, *K*_*es*_ is standard effective conductance, and *T*_*e*_ is operative temperature. Standard operative temperature equates the heat flux that an organism experiences in a black-body temperature-controlled metabolic chamber under specific convective conditions to that of a complex thermal environment to predict a physiological response. For our simulations, we assumed *K*_*es*_ represented the effective conductance at 0.1 m/s for our standard convective conditions. The specific calculations for these simulations can be found in Riddell *et al*. (2019) and the Python script can be found on GitHub (github.com/ecophysiology).

We used these simulations to isolate the thermoregulatory consequences of geographic variation in dorsal feather reflectance. We were specifically interested in estimating the water requirements for evaporative cooling (termed ‘cooling costs’) because these costs are highly relevant to the environmental pressures shaping physiological adaptation in larks (Tieleman et al. 2003*a*). We estimated cooling costs across all sites with available climate and elevational data (*n* = 66). For all sites, we generated estimates of cooling costs using two simulations: one with the average dorsal reflectance across all sites (average = 0.65) and the other incorporating the site-specific dorsal reflectance (range = 0.47 – 0.80). Ventral feather reflectance was held constant (average = 0.69) to specifically focus on the role of dorsal feather reflectance in camouflage and thermoregulation. For each site, we subtracted the cooling costs between the two simulations. The difference (termed reduction in cooling costs) provides an index for the site-specific reduction in thermoregulatory pressure driven by geographic variation in dorsal feather reflectance.

We then determined whether the reduction in cooling costs from geographic variation in reflectance was associated with climatic variables. We fit exponential models from the *nls*() function in *R* (v4.0.1; R Core Team 2020) to determine the relationship between the reduction in cooling costs and the principal components of climatic variables. For climatic variables, we averaged the principal components that described seasonality (PC1), aridity (PC2), and temperature (PC3; see Table S4 for loadings). We converted the principal components to positive values by adding the lowest value to each principal component, which generated interpretable confidence intervals. Starting values for coefficients were generated using the *nlsLM*() function. We assessed the significance of the models based on the regression coefficients, standard errors of coefficients, and 95% confidence intervals. We then used an AIC model selection framework to determine whether models with the principal components of seasonality, aridity, or temperature were more likely to explain the variation in the reduction in cooling costs attributed to variation in feather reflectance.

## Results

### Phenotypic variation

Plumage characters varied among subspecies and sexes, but also exhibited substantial overlap. The PCA of plumage characters revealed general clustering by both subspecies and sex (Figure 2A). For example, *E. a. leucansiptila* tended to have higher PC1 and PC2 scores, indicating that they were lighter and more patterned compared to other subspecies. Furthermore, males tended to have lower PC1 scores and higher PC2 scores than females, indicating that males tended to have less dorsal patterning than females on average. Comparisons of mean values using Tukey’s posthoc tests revealed various groupings among both male and female larks, but also indicated substantial overlap or gradations in phenotypes among subspecies of Horned Lark (Figure 2B–G). In parallel with PCA scores and loadings, we found that *E. a. leucansiptila* was in its own posthoc grouping for brightness for both males (Figure 2B) and females (Figure 2C). Similarly, *E. a. rubea* tended to differ from other subspecies in mean values for all three plumage variables and was frequently in its own posthoc grouping for both males and females. Box plots also revealed substantial overlap in geographically proximate subspecies. For example, the geographic distribution of *E. a. ammophila* overlaps with *E. a. actia* to the west in south-central California, and the two subspecies exhibited substantial overlap in posthoc groupings. Similar overlap was present in other pairs of geographically proximate subspecies, such as *E. a. occidentalis* and *E. a. adusta*, which come into contact in southern Arizona. These results suggest ample clinal variation among subspecies of Horned Lark, as has been noted elsewhere (Behle 1942).

**Figure 2:**
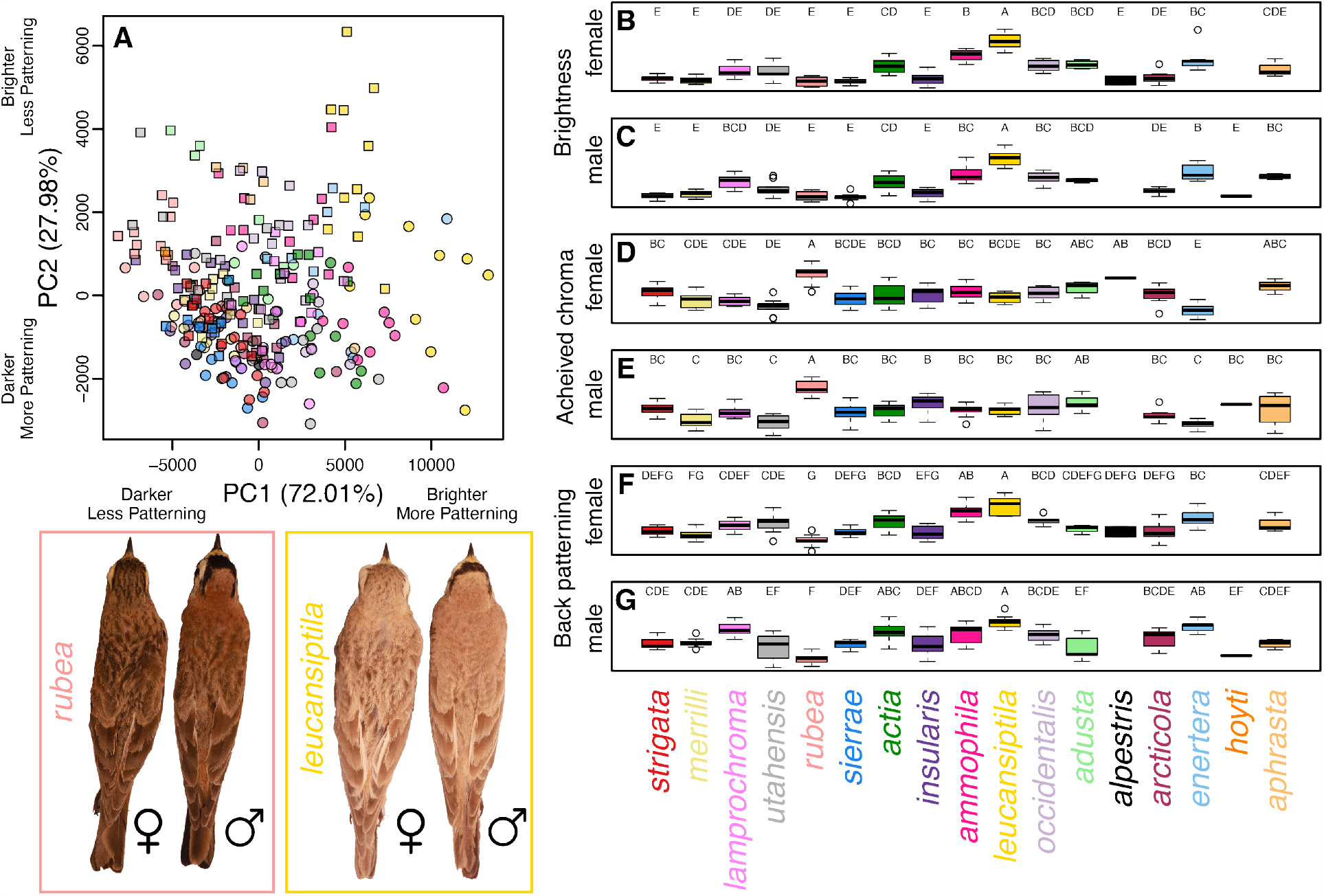
Phenotypic variation in Horned Lark plumage. A principal component analysis (A) reveals clustering by subspecies (colors correspond to labels below boxplots) and sex (males shown with squares, females shown with circles). Panels B–G show box plots of brightness, achieved chroma, and dorsal patterning for males and females. Letters above each boxplot correspond to Tukey’s posthoc groupings. Subspecies are ordered to reflect their approximate geographic distributions from northwest to southeast. Inset on the lower left shows is an example of male and female *E. a. rubea* and *E. a. leucansiptila*, which represent opposite ends of the phenotypic distribution of larks included in this study.

### Phenotype-environment associations

Using linear models, we found multiple associations between dorsal plumage variation and soil conditions underlying background matching in Horned Larks (Figure 3). Specifically, we found that dorsal brightness was positively associated with soil brightness (β_soil brightness_ = 47.17 ± 3.5; t-value = 13.49; P < 0.001) and differed marginally between sexes (β_sex_male_ = 485.46 ± 250.21; t-value =1.94; P = 0.05). Achieved chroma was positively associated with soil redness (β_soil redness_ = 2e-3 ± 3.32e-4; t-value = 6.02; P < 0.001) and differed marginally between sexes (β_sex_male_ = -0.01 ± 5.31e-3; t-value = -1.9; P = 0.06). Finally, we found that dorsal patterning was associated with soil granularity (β_soil granularity_ = 169.23 ± 19.91; t-value = 8.50; P < 0.001) such that dorsal plumages with higher contrast patterning were associated with more granular soils (i.e., more coarse fragments, more clay), and that males were less patterned than females (β_sex_male_ = -2358.47 ± 354.86; t-value = -6.65; P < 0.001).

**Figure 3:**
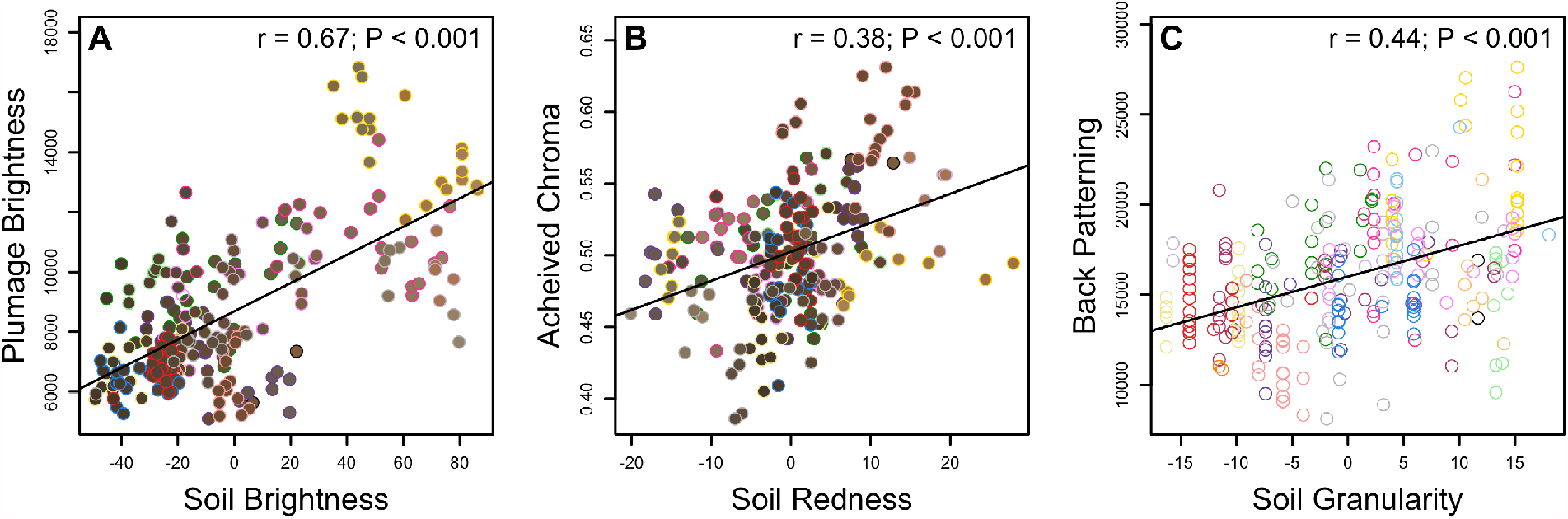
Associations between plumage and soil conditions indicating background matching in Horned Larks. In panels A and B, the fill of each point corresponds to the color of the soil for the locality of the vouchered specimen, which in turn is based on RGB values of USDA soil surveys. The outline of each point corresponds to the subspecies, as seen in Figure 1. Results from Pearson’s correlation tests are shown in the upper right hand of each plot. A trend line for the relationship between the two variables is drawn when the Pearson’s correlation test is significant (P < 0.05).

When we expanded our linear models to consider how bioclimatic and soil conditions simultaneously impact dorsal reflectance within the same model, we found a positive association between dorsal plumage brightness (UV-VIS) and soil brightness, seasonality, aridity, and temperature (Table 1). We also found positive associations between dorsal solar reflectance (UV-VIS-IR) and seasonality and aridity, but not between dorsal solar reflectance and soil brightness or temperature (Table 1).

**Table 1:** Linear model output for plumage-environment associations. *P*-values with asterisks indicate statistically significant terms (\**P* < 0.05; \*\**P* < 0.01; \*\*\**P* < 0.001).

| Response Variable | Predictor Variable | Beta + SE | t-value | *P*-value |
| --- | --- | --- | --- | --- |
| Dorsal Brightness | Intercept | 8420.53 ± 138.09 | 60.98 | <0.001*** |
|  | Sex (male) | 418.17 ± 192.97 | 2.17 | 0.03* |
|  | Soil brightness | 18.59 ± 3.58 | 5.19 | <0.001*** |
|  | BioClim PC1 (seasonality) | 0.17 ± 0.06 | 2.56 | 0.01* |
|  | BioClim PC2 (aridity) | 4.53 ± 0.38 | 11.85 | <0.001*** |
|  | BioClim PC3 (temperature) | 5.17 ± 0.92 | 5.61 | <0.001*** |
| Dorsal Solar<br>Reflectance | Intercept | 0.34 ± 6.82e-3 | 49.78 | <0.001*** |
|  | Sex (male) | 1.15e-3 ± 9.53e-3 | 0.12 | 0.90 |
|  | Soil brightness | 1.48e-4 ± 1.77e-4 | 0.84 | 0.40 |
|  | BioClim PC1 (seasonality) | 2.14e-5 ± 3.20e-6 | 6.67 | <0.001*** |
|  | BioClim PC2 (aridity) | 8.49e-5 ± 1.88e-5 | 4.50 | <0.001*** |
|  | BioClim PC3 (temperature) | 4.85e-5 ± 4.55e-5 | 1.06 | 0.29 |

**Table 2:**
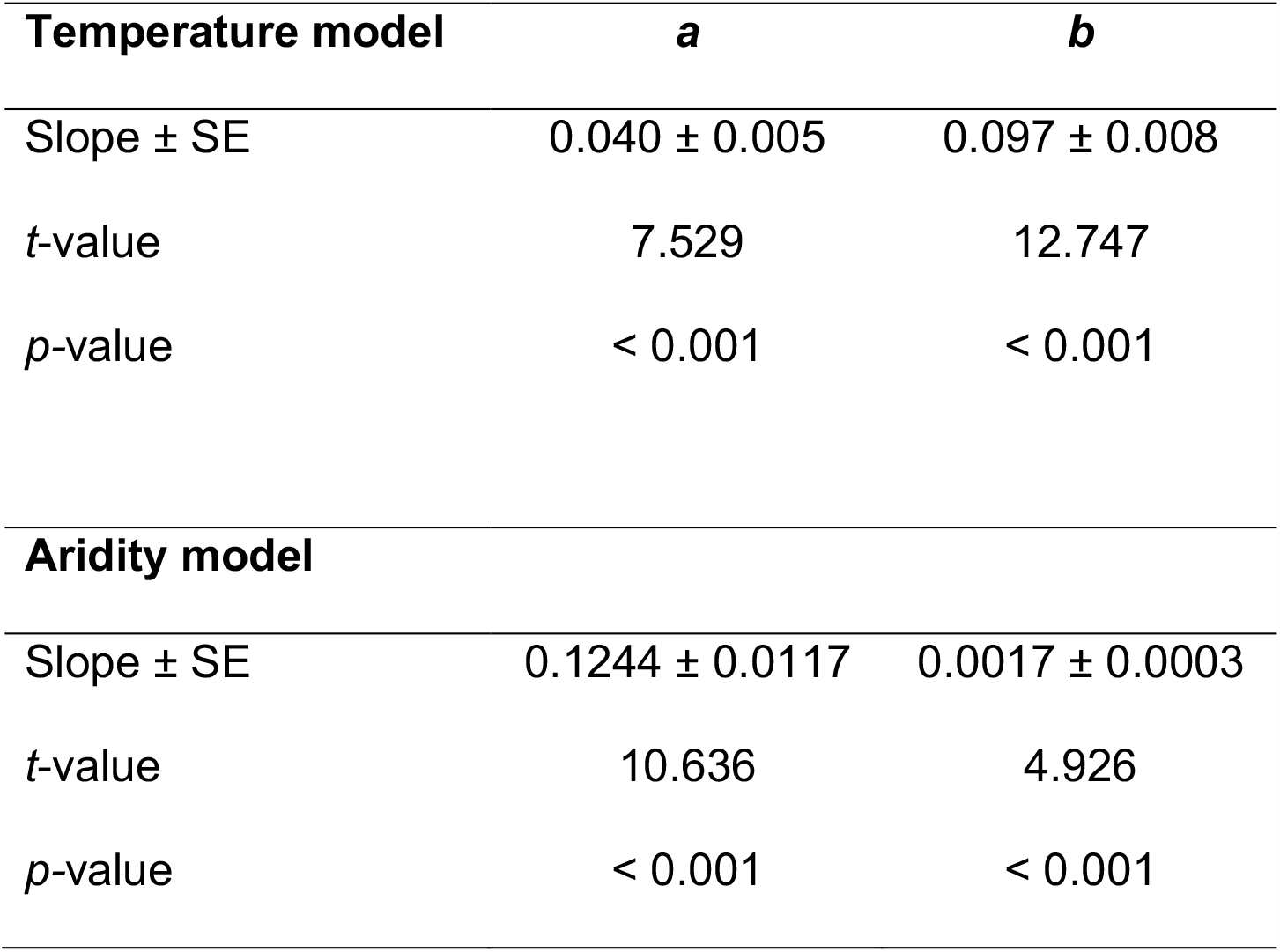
Non-linear regression analyses for investigating the relationship between the reduction in cooling costs, temperature, and aridity. Slopes and standard errors are presented and correspond to parameters in the exponential function *f*(x) = *a*^*bx*^.

| **Temperature model** | ***a*** | ***b*** |
| --- | --- | --- |
| Slope ± SE | 0.040 ± 0.005 | 0.097 ± 0.008 |
| *t*-value | 7.529 | 12.747 |
| *p*-value | < 0.001 | < 0.001 |
| **Aridity model** |  |  |
| Slope ± SE | 0.1244 ± 0.0117 | 0.0017 ± 0.0003 |
| *t*-value | 10.636 | 4.926 |
| *p*-value | < 0.001 | < 0.001 |

### Heat Flux Simulations

The observed variation in feather reflectance contributed to a substantial reduction in cooling costs compared to simulations with average feather reflectance. Incorporation of the observed variation in feather reflectance reduced water requirements for evaporative cooling by 15.1% on average (SD = 9.1%; range = 4.1% to 70.2%). The reduction in cooling costs was also positively associated with temperature (PC2) and aridity (PC3) indices but not with seasonality (PC1; Figure 4, Table S5). Although the reduction in cooling costs was associated with aridity, model selection indicated that the model with temperature far outperformed models with aridity and seasonality (Table S6).

**Figure 4:**
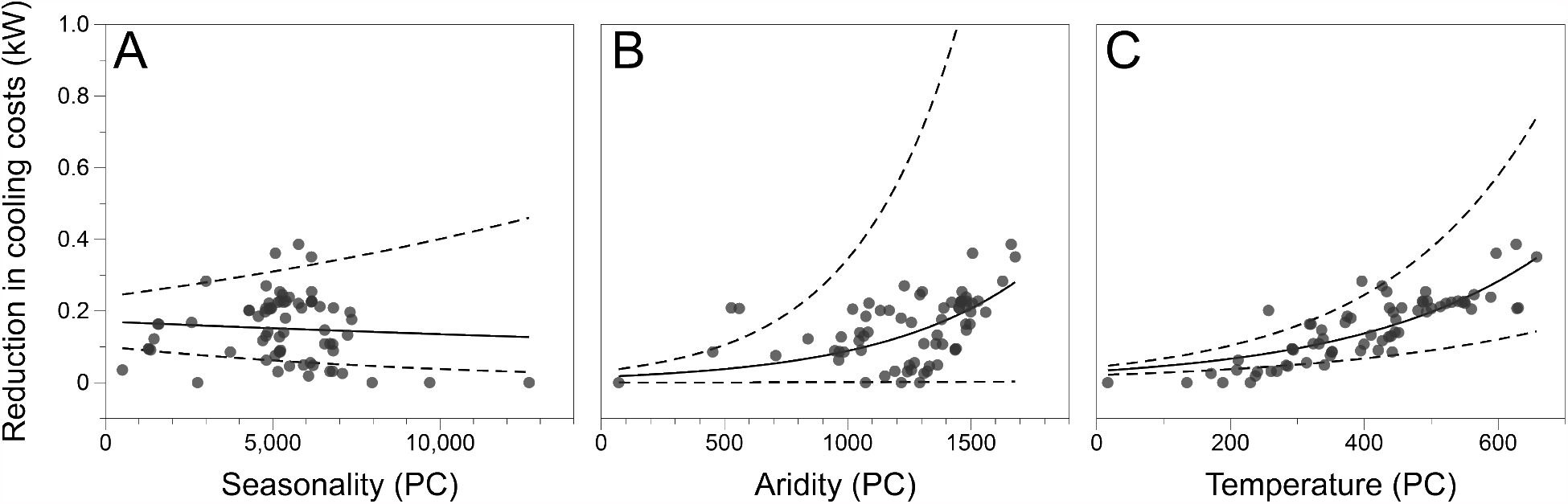
Variation in feather reflectance reduces thermoregulatory costs in hotter locations. Shown are the relationships between climatic principal components and the reduction in water requirements for evaporative cooling (termed cooling costs) due to variation in dorsal feather reflectance. The reduction in cooling costs was not associated with the principal component for (A) seasonality but was associated with (B) aridity and (C) temperature. The analyses suggest that temperature is a major driver of variation in feather reflectance for thermoregulatory purposes. Exponential models are plotted with 95% confidence intervals in the dotted line.

## Discussion

By combining data from digital photography, spectroradiometery, and remote sensing we documented multiple associations between dorsal plumage and environmental conditions of Horned Larks over a broad geographic distribution. These findings emphasize the multifarious role of feathers and animal coloration more generally. Although lark coloration has long been hypothesized to facilitate camouflage (Behle 1942; Donald et al. 2017; Mason and Unitt 2018), our study provides the first empirical evidence of phenotype-environment associations that underlie background matching in larks in brightness, hue, and patterning. Furthermore, full-spectrum spectroradiometry combined with simulation-based models of thermoregulation revealed that geographic variation in feather reflectance reduces evaporative cooling costs in hotter environments. While selective pressures imposed on total solar reflectance in feathers have just begun to be explored (Stuart-Fox et al. 2017; Medina et al. 2018), our findings suggest it plays an important role in the thermal ecology of larks.

Background matching is one component of a suite of phenotypes that organisms have evolved to avoid visual detection by predators (Merilaita and Lind 2005; Stevens and Merilaita 2009). In order to avoid such detection, natural selection has shaped the appearance of many organisms so that they resemble a random sample of the background against which their predators typically scan for prey (Endler 1978; Michalis et al. 2017). Dorsal brightness and hue in Horned Larks match the background substrate (Fig. 3A, Fig. 3B), as has also been shown in mice (Vignieri et al. 2010), gerbils (Boratyński et al. 2017), moths (Kettlewell 1955), and other taxa (Stevens et al. 2014; Troscianko et al. 2016). There is less empirical evidence for background matching in pattern as opposed to brightness or color alone, but it has been reported in cuttlefish (Barbosa et al. 2008), nightjars (Troscianko et al. 2016), and plover eggs (Stevens et al. 2017). In Horned Larks, increased dorsal ‘mottling’ with more high-contrast spots was associated with increases in sand and other coarse particles rather than with clay and silt (Fig. 3C). Beyond background matching, increased dorsal patterning in larks may also contribute to disruptive patterning that breaks up the visual outline of the bird as seen from above (Cuthill et al. 2005). Disruptive patterning in larks may be associated with molt strategies that promote the retention of worn feathers with lighter edges (Negro et al. 2019).

Our study focused on the substrate encountered by Horned Larks during the breeding months. However, migratory populations of larks must avoid detection against multiple, geographically distant substrates that differ in color and composition. Further contributing to the complexity of this challenge, soil color may change over the course of a year as precipitation increases or decreases, especially in more seasonal environments. Horned Larks molt only once per year (Pyle 1997), and thus migratory populations may need to balance competing pressures for background matching against different substrates. Future studies could expand upon our results by examining how natural selection shapes organisms with non-dynamic camouflage against multiple backgrounds.

We also uncovered associations between climate and dorsal plumage. Specifically, we found that solar reflectance (i.e., UV-VIS-NIR reflectance) is associated with two climatic variables—seasonality and aridity (Table 1)—but bears no association with soil brightness or temperature (Table 1). Thus, solar reflectance increases as climates become more arid, presumably to prevent the organism from overheating and becoming dehydrated. Interestingly, we found greater reductions in cooling costs associated with temperature than with seasonality or aridity (Fig. 4)—an observation that conflicts with implications from our linear models as we found no association between temperature and dorsal solar reflectance in our linear models (Table 1). However, the modeling approach for cooling costs incorporates additional parameters such as NIR reflectance, feather conductance, direct and diffuse solar radiation, and radiation from the ground (Gates 1980). Thus, these findings may not be in direct conflict with one another, but rather may reflect different methodologies and statistical assumptions. Regardless, NIR reflectance might play a role in allowing larks to increase solar reflectance while maintaining crypsis in the visual spectrum. Future work could further disentangle how the NIR and UV-VIS portions of light co-vary or are independently selected for optimal thermoregulation in different climates (Stuart-Fox et al. 2017).

There are many other factors beyond modifications to solar reflectance in UV-VIS and NIR wavelengths that could contribute to the ability of larks to inhabit hot, arid environments (Trost 1972; Dean and Williams 2004). First, many arid-adapted larks have reduced metabolic rates and increased water retention through various physiological adaptations (Tieleman et al. 2002). Second, behavioral adaptations also can reduce heat stress. For example, microhabitat selection such as resting in shade or animal burrows during extreme heat may contribute to thermoregulation in larks (Williams et al. 1999; Hartman and Oring 2003; Walde et al. 2009). While Horned Larks are philopatric and stay close to their breeding territory during the breeding season (Beason 1970), they are nomadic during non-breeding months and may seek out favorable habitat within reach of their individual movements to facilitate thermoregulation. Feathers are one part of a complex suite of phenotypes involved in maintaining homeostasis in thermally challenging environments. The interplay of plumage and other physiological and behavioral adaptations for thermoregulation remains an open avenue of research.

In conclusion, our study uncovered empirical evidence for the multifaceted role that plumage plays in mediating both camouflage and thermoregulation in Horned Larks. Dorsal plumage and patterning are associated with soil conditions, whereas feather solar reflectance is associated with abiotic conditions and improved cooling costs in hotter climates. Future studies could leverage these phenotype-environment associations in combination with new genomic resources (Mason et al. 2020) to identify candidate loci driving these local adaptations. Furthermore, Horned Larks are one of approximately 100 species of larks (Alaudidae) globally that vary in habitat affiliations. Phylogenetic comparative studies across the family would shed light on whether the patterns we found here are generalizable across broader taxonomic and evolutionary scales. Interactions between an organisms’ body surfaces and light from the sun are complex, and our study illustrates how natural selection has shaped the phenotypic variation across different habitats to meet potentially competing demands.

## Data and Code Availability

Raw data and code used in the analyses presented here are available via GitHub (https://github.com/mason-lab/HornedLarkCamoThermo). Raw data and metadata will be made available via Dryad upon article acceptance.

## References

Andersson, M., and L. W. Simmons. 2006. Sexual selection and mate choice. Trends in Ecology & Evolution 21:296–302.

Bakken, G. S. 1981. A two-dimensional operative-temperature model for thermal energy management by animals. Journal of Thermal Biology 6:23–30.

Barbosa, A., L. M. Mäthger, K. C. Buresch, J. Kelly, C. Chubb, C.-C. Chiao, and R. T. Hanlon. 2008. Cuttlefish camouflage: The effects of substrate contrast and size in evoking uniform, mottle or disruptive body patterns. Vision Research 48:1242–1253.

Batjes, N. H. 2016. Harmonized soil property values for broad-scale modelling (WISE30sec) with estimates of global soil carbon stocks. Geoderma 269:61–68.

Beason, R. 1970. The annual cycle of the Prairie Horned Lark in west-central Illinois (Master’s Thesis). Western Illinois University, Macomb.

Beason, R. C. 1995. Horned Lark (Eremophila alpestris), version 2.0. In The Birds of North America (A. F. Poole and F. B. Gill, Editors). Cornell Lab of Ornithology, Ithaca, NY, USA.

Beaudette, D. E., P. Roudier, and A. T. O’Geen. 2013. Algorithms for quantitative pedology: A toolkit for soil scientists. Computers & Geosciences 52:258–268.

Behle, W. 1942. Distribution and variation of the horned larks (Eremophila alpestris) of western North America. University of California Publications in Zoology 46:203–316.

Boratyński, Z., J. C. Brito, J. C. Campos, J. L. Cunha, L. Granjon, T. Mappes, A. Ndiaye, et al. 2017. Repeated evolution of camouflage in speciose desert rodents. Scientific Reports 7:1–10.

Burtt, E. H. 1981. The adaptiveness of animal colors. BioScience 31:723–729.

Burtt, E. H.. 2004. GLOGER’S RULE, FEATHER-DEGRADING BACTERIA, AND COLOR VARIATION AMONG SONG SPARROWS. Condor 106:681–686.

Campbell, G., and J. Norman. 1998. An introduction to environmental biophysics (2nd edition.). Springer-Verlag, New York, NY.

Caro, T. 2017. Wallace on Coloration: Contemporary Perspective and Unresolved Insights. Trends in Ecology & Evolution 32:23–30.

Caro, T., T. N. Sherratt, and M. Stevens. 2016. The ecology of multiple colour defences. Evolutionary Ecology 30:797–809.

Cook, L. M., and I. J. Saccheri. 2013. The peppered moth and industrial melanism: Evolution of a natural selection case study. Heredity 110:207–212.

Cott, H. B. 1944. Adaptive Coloration in Animals. Bradford and Dickens Drayton House, London.

Cuthill, I. C., W. L. Allen, K. Arbuckle, B. Caspers, G. Chaplin, M. E. Hauber, G. E. Hill, et al. 2017. The biology of color. Science 357:eaan0221.

Cuthill, I. C., M. Stevens, J. Sheppard, T. Maddocks, C. A. Párraga, and T. S. Troscianko. 2005. Disruptive coloration and background pattern matching. Nature 434:72–74.

de Mendiburu, F. 2020. agricolae: Statistical Procedures for Agricultural Research.

de Zwaan, D. R., and K. Martin. 2018. Substrate and structure of ground nests have fitness consequences for an alpine songbird. Ibis 160:790–804.

Dean, W. R. J., and J. B. Williams. 2004. Adaptations of birds for life in deserts with particular reference to Larks (Alaudidae). Transactions of the Royal Society of South Africa 59:79–91.

Delhey, K. 2019. A review of Gloger’s rule, an ecogeographical rule of colour: definitions, interpretations and evidence. Biological Reviews brv. 12503.

Delhey, K., J. Dale, M. Valcu, and B. Kempenaers. 2019. Reconciling ecogeographical rules: rainfall and temperature predict global colour variation in the largest bird radiation. (G. Grether, ed.) Ecology Letters 22:726–736.

Donald, P. F., P. Alström, and D. Engelbrecht. 2017. Possible mechanisms of substrate colour-matching in larks (Alaudidae) and their taxonomic implications. Ibis 159:699–702.

Endler, J. A. 1978. A Predator’s View of Animal Color Patterns. Evolutionary Biology 319–364.

Farkas, T. E., T. Mononen, A. A. Comeault, I. Hanski, and P. Nosil. 2013. Evolution of camouflage drives rapid ecological change in an insect community. Current Biology 23:1835–1843.

Gates, D. 1980. Solar Radiation. Biophysical Ecology, Springer Advanced Texts in Life Sciences. Springer, New York, NY.

Gloger, C. W. L. 1833. Das Abändern der Vögel durch Einfluss des Klimas. August Schulz, Breslau, Germany.

Godfrey, D., J. N. Lythgoe, and D. A. Rumball. 1987. Zebra stripes and tiger stripes: the spatial frequency distribution of the pattern compared to that of the background is significant in display and crypsis. Biological Journal of the Linnean Society 32:427–433.

Gueymard, C. A. 2001. Parameterized transmittance model for direct beam and circumsolar spectral irradiance. Solar Energy 71:325–346.

Hartman, C. A., and L. W. Oring. 2003. Orientation and microclimate of horned lark nests: the importance of shade. The Condor 105:158–163.

Hijmans, R. J., S. E. Cameron, J. L. Parra, P. G. Jones, and A. Jarvis. 2005. Very high resolution interpolated climate surfaces for global land areas. International Journal of Climatology 25:1965–1978.

Isaac, L. A., and P. T. Gregory. 2013. Can snakes hide in plain view? Chromatic and achromatic crypsis of two colour forms of the Western Terrestrial Garter Snake (Thamnophis elegans). Biological Journal of the Linnean Society 108:756–772.

Kearney, M. R., and W. P. Porter. 2020. NicheMapR – an R package for biophysical modelling: the ectotherm and Dynamic Energy Budget models. Ecography 43:85–96.

Kearney, M. R., W. P. Porter, and S. A. Murphy. 2016. An estimate of the water budget for the endangered night parrot of Australia under recent and future climates. Climate Change Responses 3:14.

Kettlewell, H. B. D. 1955. Selection experiments on industrial melanism in the Lepidoptera. Heredity 9:323–342.

Laurent, S., S. P. Pfeifer, M. L. Settles, S. S. Hunter, K. M. Hardwick, L. Ormond, V. C. Sousa, et al. 2016. The population genomics of rapid adaptation: disentangling signatures of selection and demography in white sands lizards. Molecular Ecology 25:306–323.

Linnen, C. R., E. P. Kingsley, J. D. Jensen, and H. E. Hoekstra. 2009. On the origin and spread of an adaptive allele in deer mice. Science 325:1095–1098.

Maia, R., C. M. Eliason, P.-P. Bitton, S. M. Doucet, and M. D. Shawkey. 2013. pavo : an R package for the analysis, visualization and organization of spectral data. (A. Tatem, ed.) Methods in Ecology and Evolution n/a-n/a.

Maia, R., H. Gruson, J. A. Endler, and T. E. White. 2019. PAVO 2: New tools for the spectral and spatial analysis of colour in R. (R. B. O’Hara, ed.) Methods in Ecology and Evolution 10:1097–1107.

Marcondes, R. S., K. F. Stryjewski, and R. T. Brumfield. 2020. Testing the simple and complex versions of Gloger’s rule in the Variable Antshrike (Thamnophilus caerulescens, Thamnophilidae). The Auk 137:ukaa026.

Mason, N. A., and R. C. K. Bowie. 2020. Plumage patterns: Ecological functions, evolutionary origins, and advances in quantification. The Auk ukaa060.

Mason, N. A., P. Pulgarin, C. D. Cadena, and I. J. Lovette. 2020a. De novo assembly of a high-quality reference genome for the Horned Lark (Eremophila alpestris). G3:Genes|Genomes|Genetics 10:475–478.

Mason, N. A., and P. Unitt. 2018. Rapid phenotypic change in a native bird population following conversion of the Colorado Desert to agriculture. Journal of Avian Biology 49:1–6.

Mason, N., P. Unitt, and J. Sparks. 2020b. Agriculture induces isotopic shifts and niche contraction in Horned Larks (Eremophila alpestris) of the southern Colorado Desert. Journal of Ornithology In Press.

Medina, I., E. Newton, M. R. Kearney, R. A. Mulder, W. P. Porter, and D. Stuart-Fox. 2018. Reflection of near-infrared light confers thermal protection in birds. Nature Communications 9:3610.

Merilaita, S., and J. Lind. 2005. Background-matching and disruptive coloration, and the evolution of cryptic coloration. Proceedings of the Royal Society B: Biological Sciences 272:665–670.

Merilaita, S., J. Tuomi, and V. Jormalainen. 1999. Optimization of cryptic coloration in heterogeneous habitats. Biological Journal of the Linnean Society 67:151–161.

Michalis, C., N. E. Scott-Samuel, D. P. Gibson, and I. C. Cuthill. 2017. Optimal background matching camouflage. Proceedings of the Royal Society B: Biological Sciences 284.

Nachman, M. W., H. E. Hoekstra, and S. D’Agostino. 2003. The genetic basis of adaptive melanism in pocket mice. Proceedings of the National Academy of Sciences of the United States of America 100:5268–5273.

Negro, J. J., I. Galván, and J. Potti. 2019. Adaptive plumage wear for increased crypsis in the plumage of Palearctic larks (Alaudidae). Ecology 100.

Porter, W. P., and D. M. Gates. 1969. Thermodynamic Equilibria of Animals with Environment. Ecological Monographs 39:227–244.

Pyle, P. 1997. Identification Guide to North American Birds. Part I: Columbidae to Ploceidae. Braun-Brumfield, Ann Arbor, Michigan.

R Core Team. 2020. R: A language and environment for statistical computing. R Foundation for Statistical Computing, Vienna, Austria.

Riddell, E. A., K. J. Iknayan, B. O. Wolf, B. Sinervo, and S. R. Beissinger. 2019. Cooling requirements fueled the collapse of a desert bird community from climate change. Proceedings of the National Academy of Sciences 201908791.

Rosenblum, E. B., H. Rompler, T. Schoneberg, and H. E. Hoekstra. 2009. Molecular and functional basis of phenotypic convergence in white lizards at White Sands. Proceedings of the National Academy of Sciences 107:2113–2117.

Rosenblum, E. B., H. Rompler, T. Schoneberg, and H. E. Hoekstra. 2010. Molecular and functional basis of phenotypic convergence in white lizards at White Sands. Proceedings of the National Academy of Sciences 107:2113–2117.

Sanderman, J., T. Hengl, and G. J. Fiske. 2017. Soil carbon debt of 12,000 years of human land use. Proceedings of the National Academy of Sciences 114:9575–9580.

Schneider, C. A., W. S. Rasband, and K. W. Eliceiri. 2012. NIH Image to ImageJ: 25 years of image analysis. Nature Methods 9:671–675.

Shultz, A. J., and K. J. Burns. 2017. The role of sexual and natural selection in shaping patterns of sexual dichromatism in the largest family of songbirds (Aves: Thraupidae). Evolution 71:1061–1074.

Smith, K. R., V. Cadena, J. A. Endler, M. R. Kearney, W. P. Porter, and D. Stuart-Fox. 2016. Color Change for Thermoregulation versus Camouflage in Free-Ranging Lizards. The American Naturalist 188:668–678.

Steel, R., J. Torrie, and D. Dickey. 1997. Principles and Procedures of Statistics: A Biometrical Approach. McGraw-Hill.

Stevens, M., A. E. Lown, and L. E. Wood. 2014. Color change and camouflage in juvenile shore crabs Carcinus maenas. Frontiers in Ecology and Evolution 2:1–14.

Stevens, M., and S. Merilaita. 2009. Animal camouflage: current issues and new perspectives. Philosophical Transactions of the Royal Society B: Biological Sciences 364:423–427.

Stevens, M., J. Troscianko, J. K. Wilson-Aggarwal, and C. N. Spottiswoode. 2017. Improvement of individual camouflage through background choice in ground-nesting birds. Nature Ecology & Evolution 1:1325–1333.

Stoddard, M. C., and D. Osorio. 2019. Animal coloration patterns: linking spatial vision to quantitative analysis. The American Naturalist 193:164–186.

Stoddard, M. C., and R. O. Prum. 2008. Evolution of avian plumage color in a tetrahedral color space: a phylogenetic analysis of New World buntings. The American Naturalist 171:755–776.

Stoddard, M. C., and M. Stevens. 2010. Pattern mimicry of host eggs by the Common Cuckoo, as seen through a bird’s eye. Proceedings of the Royal Society B: Biological Sciences 277:1387–1393.

Stuart-Fox, D., E. Newton, and S. Clusella-Trullas. 2017. Thermal consequences of colour and near-infrared reflectance. Philosophical Transactions of the Royal Society B: Biological Sciences 372:20160345.

Tieleman, B. I., J. B. Williams, and P. Bloomer. 2003a. Adaptation of metabolism and evaporative water loss along an aridity gradient. Proceedings of the Royal Society of London. Series B: Biological Sciences 270:207–214.

Tieleman, B. I., J. B. Williams, and M. E. Buschur. 2002. Physiological Adjustments to Arid and Mesic Environments in Larks (Alaudidae). Physiological and Biochemical Zoology 75:305–313.

Tieleman, B. I., J. B. Williams, M. E. Buschur, and C. R. Brown. 2003b. PHENOTYPIC VARIATION OF LARKS ALONG AN ARIDITY GRADIENT: ARE DESERT BIRDS MORE FLEXIBLE? Ecology 84:1800–1815.

Troscianko, J., and M. Stevens. 2015. Image calibration and analysis toolbox - a free software suite for objectively measuring reflectance, colour and pattern. (S. Rands, ed.) Methods in Ecology and Evolution 6:1320–1331.

Troscianko, J., J. Wilson-Aggarwal, M. Stevens, and C. N. Spottiswoode. 2016. Camouflage predicts survival in ground-nesting birds. Scientific Reports 6:19966.

Trost, C. H. 1972. Adaptations of Horned Larks (Eremophila alpestris) to Hot Environments. The Auk 89:506–527.

Van’t Hof, A. E., N. Edmonds, M. Dalíková, F. Marec, and I. J. Saccheri. 2011. Industrial melanism in british peppered moths has a singular and recent mutational origin. Science 332:958–960.

Vignieri, S. N., J. G. Larson, and H. E. Hoekstra. 2010. The selective advantage of crypsis in mice. Evolution 64:2153–2158.

Vo, A.-T. E., M. S. Bank, J. P. Shine, and S. V. Edwards. 2011. Temporal increase in organic mercury in an endangered pelagic seabird assessed by century-old museum specimens. Proceedings of the National Academy of Sciences 108:7466–7471.

Vorobyev, M., and D. Osorio. 1998. Receptor noise as a determinant of colour thresholds. Proceedings of the Royal Society of London. Series B: Biological Sciences 265:351–358.

Vorobyev, M., D. Osorio, A. T. D. Bennett, N. J. Marshall, and I. C. Cuthill. 1998. Tetrachromacy, oil droplets and bird plumage colours. Journal of Comparative Physiology A: Sensory, Neural, and Behavioral Physiology 183:621–633.

Wacker, C. B., B. M. McAllan, G. Körtner, and F. Geiser. 2016. The functional requirements of mammalian hair: a compromise between crypsis and thermoregulation? The Science of Nature 103:53.

Walde, A. D., A. M. Walde, D. K. Delaney, and L. L. Pater. 2009. Burrows of Desert Tortoises (Gopherus agassizii) as Thermal Refugia for Horned Larks (Eremophila alpestris) in the Mojave Desert. The Southwestern Naturalist 54:375–381.

Walsberg, G. E. 1983. Coat Color and Solar Heat Gain in Animals. BioScience 33:88–91.

Walsberg, G. E. 1988. Heat flow through avian plumages: The relative importance of conduction, convection, and radiation. Journal of Thermal Biology 13:89–92.

Williams, J. B., B. I. Tieleman, and M. Shobrak. 1999. Lizard Burrows Provide Thermal Refugia for Larks in the Arabian Desert. The Condor 101:714–717.

Wolf, B. O., and G. E. Walsberg. 2000. The Role of the Plumage in Heat Transfer Processes of Birds. American Zoologist 40:575–584.

Zink, R. M., and J. V. Remsen. 1986. Evolutionary Processes and Patterns of Geographic Variation in Birds. Current Ornithology 4:1–69.

