## Supplementary Material for "Plumage balances camouflage and thermoregulation in Horned Larks (*Eremophila alpestris*)"

The authors wish to be identified to the reviewers.


Nicholas A. Mason[1,2], Eric A. Riddell[1,3], Felisha Romero[1], Carla Cicero[1], Rauri C.K. Bowie[1,4]

[1]Museum of Vertebrate Zoology, University of California, Berkeley, California, USA

[2]Museum of Natural Science and Department of Biological Sciences, Louisiana State University, Baton Rouge, Louisiana, USA

[3]Iowa State University, Department of Ecology, Evolution, and Organismal Biology, Ames, Iowa, USA

[4]Department of Integrative Biology, University of California, Berkeley, California, USA

Supplementary Information

Supplementary Figure S1: Example of photography set up and dorsal region of interest (back).

Supplementary Table S1: Voucher information for 270 individuals of Horned Lark (*Eremophila alpestris*) included in this study.

Supplementary Table S2: Principal component analysis loadings for plumage data set.

Supplementary Table S3: Principal component analysis loadings for soil color data set.

Supplementary Table S4: Principal component analysis loadings for soil granularity data set.

Supplementary Table S5: Principal component analysis loadings for WorldClim bioclimatic data set.

Supplementary Table S6: Model selection for climatic covariates and reduction in cooling costs.

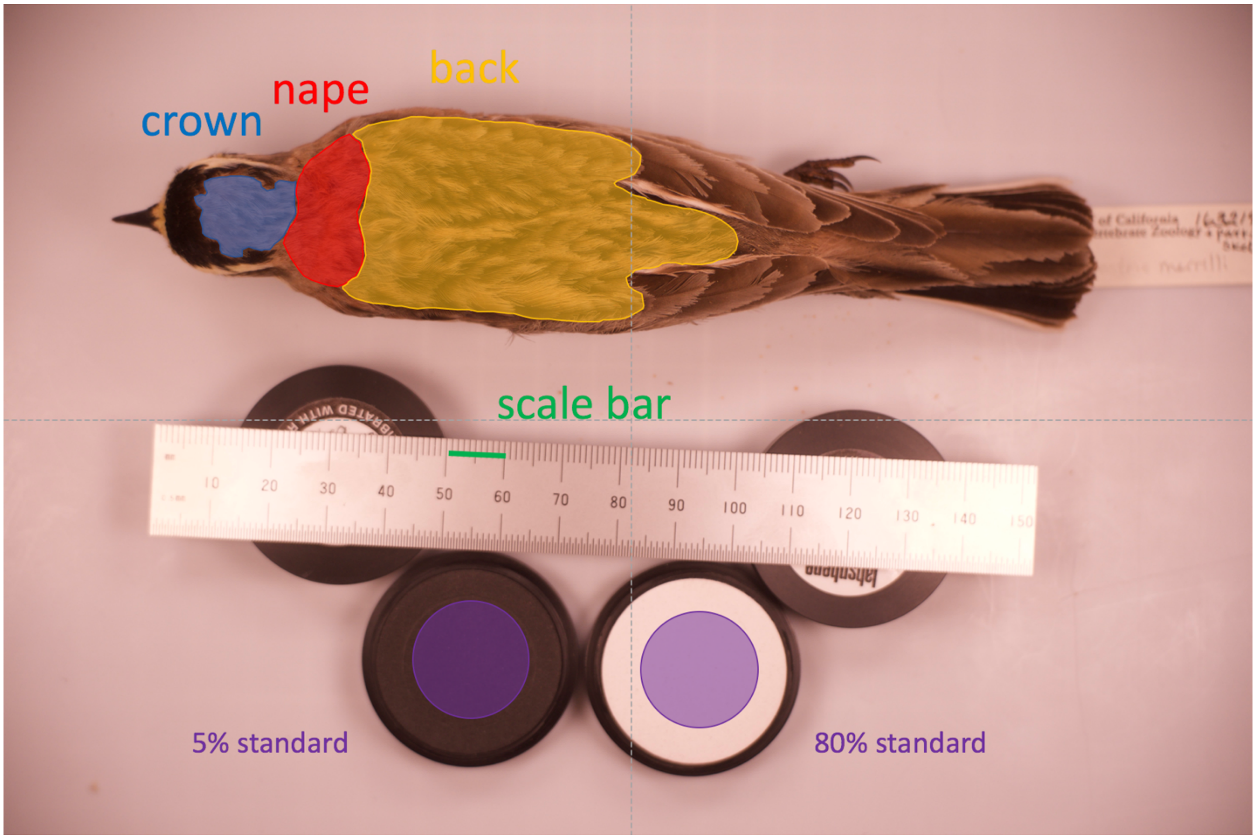


Supplementary Figure S1: Photography setup and example of polygon corresponding to region of interest (back) measured among Horned Larks (*Eremophila alpestris*) in this study.

Supplementary Table S1: Voucher information for 270 individuals of Horned Lark (*Eremophila alpestris*) included in this study.

| Subspecies | MVZ catalog number | Sex | Year | Month | Day | Country | State / Province | Locality | Longitude | Latitude |
|---|---|---|---|---|---|---|---|---|---|---|
| *actia* | 3667 | male | 1908 | 05 | 26 | USA | California | Chula Vista | 32.64 | -117.083 |
| *actia* | 3471 | female | 1908 | 04 | 06 | USA | California | Nat'l City | 32.675 | -117.109 |
| *actia* | 3472 | female | 1908 | 04 | 06 | USA | California | Nat'l City | 32.675 | -117.109 |
| *actia* | 3473 | female | 1908 | 04 | 06 | USA | California | Nat'l City | 32.675 | -117.109 |
| *actia* | 3479 | male | 1908 | 04 | 10 | USA | California | False Bay | 32.808 | -117.267 |
| *actia* | 19488 | male | 1911 | 04 | 30 | USA | California | Earlimart | 35.882 | -119.271 |
| *actia* | 19496 | female | 1911 | 05 | 03 | USA | California | Earlimart | 35.882 | -119.271 |
| *actia* | 19486 | male | 1911 | 04 | 24 | USA | California | Tipton | 36.06 | -119.311 |
| *actia* | 19487 | female | 1911 | 04 | 24 | USA | California | Tipton | 36.06 | -119.311 |
| *actia* | 29155 | female | 1918 | 06 | 27 | USA | California | """Hayes Station"", 19 mi SW Mendota" | 36.676 | -120.611 |
| *actia* | 64599 | female | 1934 | 03 | 19 | USA | California | 2 mi SW Friant | 36.965 | -119.728 |
| *actia* | 90619 | male | 1941 | 05 | 19 | USA | California | Santa Cruz | 36.975 | -122.007 |
| *actia* | 120329 | female | 1947 | 03 | 07 | USA | California | Santa Cruz | 36.986 | -122.02 |
| *actia* | 120330 | male | 1947 | 03 | 07 | USA | California | Santa Cruz | 36.986 | -122.02 |
| *actia* | 19481 | male | 1911 | 03 | 21 | USA | California | Los Banos | 37.061 | -120.848 |
| *actia* | 19482 | female | 1911 | 03 | 23 | USA | California | Los Banos | 37.061 | -120.848 |
| *actia* | 19483 | male | 1911 | 03 | 23 | USA | California | Los Banos | 37.061 | -120.848 |
| *actia* | 19484 | male | 1911 | 03 | 23 | USA | California | Los Banos | 37.061 | -120.848 |

| | | | | | | | | | | |
|---|---|---|---|---|---|---|---|---|---|---|
| *actia* | 116808 | male | 1897 | 03 | 22 | USA | California | San Jose | 37.324 | -121.89 |
| *actia* | 116809 | female | 1897 | 03 | 22 | USA | California | San Jose | 37.324 | -121.89 |
| *adusta* | 10234 | male | 1896 | 05 | 24 | USA | Arizona | Huachuca Mts. | 31.489 | -110.407 |
| *adusta* | 10235 | female | 1896 | 06 | 30 | USA | Arizona | Huachuca Mts. | 31.489 | -110.407 |
| *adusta* | 24483 | male | 1913 | 10 | 05 | USA | Arizona | Fort Huachuca | 31.553 | -110.347 |
| *adusta* | 5433 | male | 1885 | 02 | 03 | USA | Arizona | Fort Huachuca | 31.553 | -110.347 |
| *adusta* | 5434 | male | 1885 | 02 | 11 | USA | Arizona | Fort Huachuca | 31.553 | -110.347 |
| *adusta* | 5435 | female | 1885 | 02 | 03 | USA | Arizona | Fort Huachuca | 31.553 | -110.347 |
| *adusta* | 5436 | female | 1885 | 02 | 03 | USA | Arizona | Fort Huachuca | 31.553 | -110.347 |
| *adusta* | 78852 | male | 1894 | 05 | 24 | USA | Arizona | Willcox | 32.253 | -109.831 |
| *adusta* | 58954 | female | 1931 | 08 | 30 | USA | New Mexico | Horse Camp, Victoria Cattle Co., Animas Valley | 31.612 | -108.871 |
| *alpestris* | 63384 | female | 1875 | 03 | 27 | USA | Massachusetts | Swampscott | 42.47 | -70.91 |
| *alpestris* | 63386 | female | 1878 | 03 | 16 | USA | Massachusetts | Swampscott | 42.47 | -70.91 |
| *ammophila* | 140817 | female | 1960 | 04 | 12 | USA | California | Pinto Basin | 33.936 | -115.69 |
| *ammophila* | 24550 | male | 1914 | 03 | 20 | USA | California | Victorville, Mohave River | 34.536 | -117.29 |
| *ammophila* | 29166 | male | 1918 | 03 | 14 | USA | California | Mohvave | 35.053 | -118.171 |
| *ammophila* | 29169 | female | 1918 | 03 | 14 | USA | California | Mohave | 35.053 | -118.171 |
| *ammophila* | 29170 | female | 1918 | 03 | 14 | USA | California | Mohave | 35.053 | -118.171 |
| *ammophila* | 29172 | male | 1918 | 03 | 15 | USA | California | Mohave | 35.053 | -118.171 |
| *ammophila* | 29173 | male | 1918 | 03 | 15 | USA | California | Mohave | 35.053 | -118.171 |
| *ammophila* | 29177 | female | 1918 | 03 | 15 | USA | California | Mojave | 35.053 | -118.171 |
| *ammophila* | 82393 | female | 1923 | 04 | 23 | USA | California | Taft | 35.142 | -119.456 |
| *ammophila* | 82394 | male | 1923 | 04 | 24 | USA | California | Taft | 35.146 | -119.456 |
| *ammophila* | 102726 | female | 1923 | 05 | 06 | USA | California | Buena Vista Lake | 35.188 | -119.287 |

| | | | | | | | | | | |
|---|---|---|---|---|---|---|---|---|---|---|
| *ammophila* | 102727 | male | 1923 | 04 | 29 | USA | California | Buena Vista Lake | 35.194 | -119.3 |
| *ammophila* | 19502 | female | 1911 | 05 | 23 | USA | California | 7 mi SE Simmler, Carrizo Plains | 35.279 | -119.902 |
| *ammophila* | 29153 | male | 1918 | 04 | 17 | USA | California | 2 mi N Edison Station | 35.375 | -118.878 |
| *ammophila* | 22501 | male | 1912 | 04 | 28 | USA | California | Keeler | 36.489 | -117.87 |
| *ammophila* | 22502 | female | 1912 | 04 | 28 | USA | California | Keeler | 36.489 | -117.87 |
| *ammophila* | 162726 | male | 1971 | 05 | 22 | USA | Nevada | Send Sarcobatus Flat, 7 mi W, 2 mi S Springdale | 37.001 | -116.881 |
| *ammophila* | 162727 | male | 1971 | 05 | 22 | USA | Nevada | Send Sarcobatus Flat, 7 mi W, 2 mi S Springdale | 37.001 | -116.881 |
| *ammophila* | 162729 | female | 1971 | 05 | 22 | USA | Nevada | Send Sarcobatus Flat, 7 mi W, 2 mi S Springdale | 37.001 | -116.881 |
| *ammophila* | 162730 | female | 1971 | 05 | 22 | USA | Nevada | Send Sarcobatus Flat, 7 mi W, 2 mi S Springdale | 37.001 | -116.881 |
| *aphrasta* | 152933 | male | 1964 | 05 | 02 | Mexico | Chihuahua | Ojo Laguna | 29.462 | -106.33 |
| *aphrasta* | 152934 | female | 1964 | 05 | 02 | Mexico | Chihuahua | Ojo Laguna | 29.462 | -106.33 |
| *aphrasta* | 143776 | male | 1961 | 09 | 02 | Mexico | Chihuahua | Ojo de Laguna, 25 mi S Gallego | 29.48 | -106.402 |
| *aphrasta* | 139540 | female | 1959 | 06 | 21 | Mexico | Chihuahua | Sierra del Nido, Canon del Alamo | 29.485 | -106.77 |
| *aphrasta* | 139544 | male | 1959 | 06 | 21 | Mexico | Chihuahua | Sierra del Nido, Canon del Alamo | 29.485 | -106.77 |
| *aphrasta* | 139545 | male | 1959 | 06 | 21 | Mexico | Chihuahua | Sierra del Nido, Canon del Alamo | 29.485 | -106.77 |
| *aphrasta* | 139546 | female | 1959 | 06 | 21 | Mexico | Chihuahua | Sierra del Nido, Canon del Alamo | 29.485 | -106.77 |
| *arcticola* | 158332 | male | 1960 | 05 | 22 | USA | Alaska | 9 mi ESE Cape Thompson | 68.087 | -165.578 |
| *arcticola* | 158334 | female | 1960 | 05 | 29 | USA | Alaska | 9 mi ESE Cape Thompson | 68.087 | -165.578 |
| *arcticola* | 158335 | male | 1960 | 05 | 29 | USA | Alaska | 9 mi ESE Cape Thompson | 68.087 | -165.578 |
| *arcticola* | 158337 | female | 1960 | 05 | 29 | USA | Alaska | 9 mi ESE Cape Thompson | 68.087 | -165.578 |
| *arcticola* | 158333 | male | 1961 | 05 | 28 | USA | Alaska | 6 mi ESE Cape Thompson | 68.1 | -165.767 |

| | | | | | | | | | | |
|---|---|---|---|---|---|---|---|---|---|---|
| *arcticola* | 102623 | male | 1888 | 04 | 17 | Canada | British Columbia | Chilliwack | 49.175 | -121.942 |
| *arcticola* | 102618 | female | 1919 | 08 | 28 | Canada | British Columbia | Pearson Mt., Headwaters Keremeos Cr. | 49.179 | -119.776 |
| *arcticola* | 102612 | female | 1925 | 10 | 14 | Canada | British Columbia | Vancouver Island, Comox | 49.683 | -124.933 |
| *arcticola* | 102614 | female | 1925 | 12 | 17 | Canada | British Columbia | Vancouver Island, Comox | 49.683 | -124.933 |
| *arcticola* | 102604 | male | 1930 | 03 | 30 | Canada | British Columbia | Okanagan Landing | 50.233 | -119.35 |
| *arcticola* | 102608 | male | 1929 | 09 | 07 | Canada | British Columbia | Okanagan Landing | 50.233 | -119.35 |
| *arcticola* | 102610 | female | 1929 | 09 | 25 | Canada | British Columbia | Okanagan Landing | 50.233 | -119.35 |
| *arcticola* | 42658 | female | 1921 | 10 | 04 | Canada | British Columbia | Okanagan Landing | 50.233 | -119.35 |
| *arcticola* | 42659 | female | 1921 | 10 | 04 | Canada | British Columbia | Okanagan Landing | 50.233 | -119.35 |
| *arcticola* | 82368 | female | 1913 | 09 | 30 | Canada | British Columbia | Okanagan Landing | 50.233 | -119.35 |
| *arcticola* | 102600 | female | 1920 | 05 | 15 | Canada | British Columbia | Masset, Graham Island, Queen Charlotte Islands | 54.017 | -132.15 |
| *arcticola* | 102601 | male | 1920 | 05 | 15 | Canada | British Columbia | Masset, Graham Island, Queen Charlotte Islands | 54.017 | -132.15 |
| *arcticola* | 102593 | male | 1924 | 06 | 19 | Canada | British Columbia | Atlin | 59.567 | -133.7 |
| *arcticola* | 102594 | male | 1924 | 07 | 08 | Canada | British Columbia | Atlin | 59.567 | -133.7 |
| *arcticola* | 102596 | female | 1924 | 08 | 07 | Canada | British Columbia | Atlin | 59.567 | -133.7 |
| *enertera* | 59518 | male | 1931 | 01 | 09 | Mexico | Baja California | San Augustin | 29.922 | -114.969 |
| *enertera* | 59563 | female | 1931 | 06 | 03 | Mexico | Baja California Sur | Magdalena, Santa Magdalena Id. | 24.909 | -112.222 |
| *enertera* | 59534 | female | 1931 | 05 | 13 | Mexico | Baja California Sur | Hiray, Llano de Hiray | 25.344 | -111.938 |
| *enertera* | 59535 | male | 1931 | 05 | 13 | Mexico | Baja California Sur | Hiray, Llano de Hiray | 25.344 | -111.938 |
| *enertera* | 59538 | female | 1931 | 05 | 13 | Mexico | Baja California Sur | Hiray, Llano de Hiray | 25.344 | -111.938 |
| *enertera* | 59539 | female | 1931 | 05 | 13 | Mexico | Baja California Sur | Hiray, Llano de Hiray | 25.344 | -111.938 |
| *enertera* | 59541 | male | 1931 | 05 | 13 | Mexico | Baja California Sur | Hiray, Llano de Hiray | 25.344 | -111.938 |
| *enertera* | 59542 | male | 1931 | 05 | 14 | Mexico | Baja California Sur | Hiray, Llano de Hiray | 25.344 | -111.938 |

| | | | | | | | | | | |
|---|---|---|---|---|---|---|---|---|---|---|
| *enertera* | 59543 | female | 1931 | 05 | 14 | Mexico | Baja California Sur | Hiray, Llano de Hiray | 25.344 | -111.938 |
| *enertera* | 59545 | female | 1931 | 05 | 14 | Mexico | Baja California Sur | Hiray, Llano de Hiray | 25.344 | -111.938 |
| *enertera* | 59552 | male | 1931 | 05 | 14 | Mexico | Baja California Sur | Hiray, Llano de Hiray | 25.344 | -111.938 |
| *enertera* | 59558 | male | 1931 | 05 | 15 | Mexico | Baja California Sur | Hiray, Llano de Hiray | 25.344 | -111.938 |
| *enertera* | 59559 | male | 1931 | 05 | 15 | Mexico | Baja California Sur | Hiray, Llano de Hiray | 25.344 | -111.938 |
| *enertera* | 59560 | female | 1931 | 05 | 15 | Mexico | Baja California Sur | Hiray, Llano de Hiray | 25.344 | -111.938 |
| *hoyti* | 82371 | male | 1922 | 03 | 14 | Canada | British Columbia | Okanagan Landing | 50.233 | -119.35 |
| *hoyti* | 73374 | male | 1936 | 06 | 23 | Canada | Manitoba | Churchill | 58.769 | -94.165 |
| *insularis* | 177000 | male | 1995 | 04 | 13 | USA | California | Santa Catalina Id., 1.25 mi N Little Harbor | 33.403 | -118.477 |
| *insularis* | 177001 | male | 1995 | 04 | 13 | USA | California | Santa Catalina Id., 1.25 mi N Little Harbor | 33.403 | -118.477 |
| *insularis* | 177002 | female | 1995 | 04 | 13 | USA | California | Santa Catalina Id., 1.25 mi N Little Harbor | 33.403 | -118.477 |
| *insularis* | 177003 | female | 1995 | 04 | 13 | USA | California | Santa Catalina Id., 1.25 mi N Little Harbor | 33.403 | -118.477 |
| *insularis* | 177004 | male | 1995 | 04 | 13 | USA | California | Santa Catalina Id., 1.25 mi N Little Harbor | 33.403 | -118.477 |
| *insularis* | 177005 | male | 1995 | 04 | 13 | USA | California | Santa Catalina Id., 1.25 mi N Little Harbor | 33.403 | -118.477 |
| *insularis* | 177006 | female | 1995 | 04 | 13 | USA | California | Santa Catalina Id., 1.25 mi N Little Harbor | 33.403 | -118.477 |
| *insularis* | 176999 | female | 1995 | 04 | 14 | USA | California | Santa Catalina Id., 2.5 mi N Little Harbor | 33.42 | -118.478 |
| *insularis* | 177009 | male | 1995 | 04 | 14 | USA | California | W end Santa Catalina Id., 1 mi S and 2 mi E Silver Pk. | 33.445 | -118.533 |
| *insularis* | 177010 | female | 1995 | 04 | 14 | USA | California | W end Santa Catalina Id., 1 mi S and 2 mi E Silver Pk. | 33.445 | -118.533 |
| *insularis* | 174005 | male | 1994 | 04 | 30 | USA | California | W end Santa Cruz Id., Canada de Los Sauces | 34.008 | -119.848 |
| *insularis* | 174006 | male | 1994 | 04 | 30 | USA | California | W end Santa Cruz Id., Canada de Los Sauces | 34.008 | -119.848 |
| *insularis* | 58402 | female | 1931 | 05 | 23 | USA | California | S slope Mt.Diablo, Santa Cruz Island | 34.026 | -119.783 |

| | | | | | | | | | | |
|---|---|---|---|---|---|---|---|---|---|---|
| *insularis* | 120124 | female | 1950 | 03 | 07 | USA | California | Forney Cove, Santa Cruz Id. | 34.059 | -119.918 |
| *insularis* | 120125 | male | 1950 | 03 | 07 | USA | California | Forney Cove, Santa Cruz Id. | 34.059 | -119.918 |
| *insularis* | 120127 | male | 1950 | 03 | 07 | USA | California | Forney Cove, Santa Cruz Id. | 34.059 | -119.918 |
| *insularis* | 120131 | female | 1950 | 03 | 08 | USA | California | Forney Cove, Santa Cruz Id. | 34.059 | -119.918 |
| *insularis* | 120132 | female | 1950 | 03 | 08 | USA | California | Forney Cove, Santa Cruz Id. | 34.059 | -119.918 |
| *insularis* | 120133 | female | 1950 | 03 | 08 | USA | California | Forney Cove, Santa Cruz Id. | 34.059 | -119.918 |
| *insularis* | 174004 | male | 1994 | 04 | 30 | USA | California | W end Santa Cruz Id., Canada de Los Sauces | 34.066 | -119.907 |
| *lamprochroma* | 132294 | female | 1954 | 05 | 31 | USA | California | Cabin Creek, White Mts. | 37.713 | -118.261 |
| *lamprochroma* | 132295 | male | 1954 | 06 | 02 | USA | California | Cabin Creek, White Mts. | 37.713 | -118.261 |
| *lamprochroma* | 165717 | female | 1978 | 06 | 21 | USA | Nevada | Clayton Valley, 7 mi S and 2 mi W Silver Pk. | 37.744 | -117.59 |
| *lamprochroma* | 165718 | female | 1978 | 06 | 21 | USA | Nevada | Clayton Valley, 7 mi S and 2 mi W Silver Pk. | 37.744 | -117.59 |
| *lamprochroma* | 165720 | male | 1978 | 06 | 21 | USA | Nevada | Clayton Valley, 7 mi S and 2 mi W Silver Pk. | 37.744 | -117.59 |
| *lamprochroma* | 165729 | male | 1978 | 06 | 22 | USA | Nevada | Clayton Valley, 7 mi S and 2 mi W Silver Pk. | 37.744 | -117.59 |
| *lamprochroma* | 165730 | female | 1978 | 06 | 22 | USA | Nevada | Clayton Valley, 7 mi S and 2 mi W Silver Pk. | 37.744 | -117.59 |
| *lamprochroma* | 165396 | female | 1977 | 06 | 28 | USA | Nevada | Bald Mt. N slope | 38.534 | -119.115 |
| *lamprochroma* | 165398 | male | 1977 | 06 | 29 | USA | Nevada | Bald Mt. N slope | 38.534 | -119.115 |
| *lamprochroma* | 165400 | female | 1977 | 06 | 29 | USA | Nevada | Bald Mt. N slope | 38.534 | -119.115 |
| *lamprochroma* | 165401 | female | 1977 | 06 | 29 | USA | Nevada | Bald Mt. N slope | 38.534 | -119.115 |
| *lamprochroma* | 165402 | male | 1977 | 06 | 29 | USA | Nevada | Bald Mt. N slope | 38.534 | -119.115 |

| | | | | | | | | | | |
|---|---|---|---|---|---|---|---|---|---|---|
| *lamprochroma* | 162686 | male | 1971 | 06 | 07 | USA | Nevada | 5.5 mi S and 6 mi E Wellington | 38.677 | -119.264 |
| *lamprochroma* | 162687 | female | 1971 | 06 | 07 | USA | Nevada | 5.5 mi S and 6 mi E Wellington | 38.677 | -119.264 |
| *lamprochroma* | 162689 | male | 1971 | 06 | 07 | USA | Nevada | 5.5 mi S and 6 mi E Wellington | 38.677 | -119.264 |
| *lamprochroma* | 162691 | male | 1971 | 06 | 08 | USA | Nevada | 2 mi N Lobdell summit, Pine Grove Hills | 38.68 | -119.17 |
| *lamprochroma* | 180084 | male | 2002 | 05 | 30 | USA | Nevada | 4 mi N and 2 mi E Poodle Mtn., 6000 ft., Buffalo Hills, Washoe Co., Nev. | 40.876 | -119.617 |
| *lamprochroma* | 180088 | male | 2002 | 05 | 30 | USA | Nevada | 4 mi N and 2 mi E Poodle Mtn., 6000 ft., Buffalo Hills, Washoe Co., Nev. | 40.876 | -119.617 |
| *lamprochroma* | 180089 | female | 2002 | 05 | 30 | USA | Nevada | 4 mi N and 2 mi E Poodle Mtn., 6000 ft., Buffalo Hills, Washoe Co., Nev. | 40.876 | -119.617 |
| *lamprochroma* | 180090 | female | 2002 | 05 | 30 | USA | Nevada | 4 mi N and 2 mi E Poodle Mtn., 6000 ft., Buffalo Hills, Washoe Co., Nev. | 40.876 | -119.617 |
| *leucansiptila* | 8090 | female | 1909 | 03 | 30 | USA | California | Coyote Well | 32.735 | -115.967 |
| *leucansiptila* | 102733 | female | 1916 | 03 | 21 | USA | California | near Sand Dunes | 33.059 | -115.224 |
| *leucansiptila* | 102734 | female | 1916 | 03 | 21 | USA | California | near Sand Dunes | 33.059 | -115.224 |
| *leucansiptila* | 1066 | female | 1908 | 04 | 28 | USA | California | Mecca | 33.572 | -116.073 |
| *leucansiptila* | 854 | male | 1908 | 04 | 15 | USA | California | Mecca | 33.572 | -116.073 |
| *leucansiptila* | 961 | male | 1908 | 04 | 17 | USA | California | Mecca | 33.572 | -116.073 |
| *leucansiptila* | 852 | male | 1908 | 04 | 10 | USA | California | Mecca | 33.573 | -116.067 |
| *leucansiptila* | 853 | male | 1908 | 04 | 15 | USA | California | Mecca | 33.573 | -116.067 |
| *leucansiptila* | 855 | male | 1908 | 04 | 15 | USA | California | Mecca | 33.573 | -116.067 |
| *leucansiptila* | 966 | female | 1908 | 04 | 24 | USA | California | Mecca | 33.573 | -116.067 |
| *leucansiptila* | 74252 | female | 1938 | 05 | 27 | USA | California | Providence Mts., head of Cedar Canyon | 35.176 | -115.398 |
| *leucansiptila* | 74253 | female | 1938 | 05 | 27 | USA | California | Providence Mts., head of Cedar Canyon | 35.176 | -115.398 |
| *leucansiptila* | 74254 | female | 1938 | 05 | 27 | USA | California | Providence Mts., head of Cedar Canyon | 35.176 | -115.398 |

| | | | | | | | | | | |
|---|---|---|---|---|---|---|---|---|---|---|
| *leucansiptila* | 74255 | male | 1938 | 05 | 27 | USA | California | Providence Mts., head of Cedar Canyon | 35.176 | -115.398 |
| *leucansiptila* | 74457 | male | 1938 | 05 | 27 | USA | California | Providence Mts., head of Cedar Canyon | 35.176 | -115.398 |
| *leucansiptila* | 74245 | male | 1938 | 05 | 14 | USA | California | 1 mi E Cima | 35.238 | -115.481 |
| *leucansiptila* | 74248 | male | 1938 | 05 | 16 | USA | California | 3/4 mi E Cima | 35.238 | -115.485 |
| *leucansiptila* | 74249 | male | 1938 | 05 | 16 | USA | California | 3/4 mi E Cima | 35.238 | -115.485 |
| *leucansiptila* | 74244 | female | 1938 | 05 | 12 | USA | California | Cima | 35.238 | -115.498 |
| *leucansiptila* | 74251 | female | 1938 | 05 | 18 | USA | California | 2 mi NNE Cima | 35.265 | -115.498 |
| *merrilli* | 163234 | male | 1972 | 03 | 30 | USA | California | Gazelle 2 mi NW, Cleland Ranch | 41.542 | -122.546 |
| *merrilli* | 163237 | male | 1972 | 03 | 30 | USA | California | Gazelle 2 mi NW, Cleland Ranch | 41.542 | -122.546 |
| *merrilli* | 163239 | female | 1972 | 03 | 30 | USA | California | Gazelle 2 mi NW, Cleland Ranch | 41.542 | -122.546 |
| *merrilli* | 163242 | male | 1972 | 03 | 30 | USA | California | Gazelle 2 mi NW, Cleland Ranch | 41.542 | -122.546 |
| *merrilli* | 163243 | male | 1972 | 03 | 30 | USA | California | Gazelle 2 mi NW, Cleland Ranch | 41.542 | -122.546 |
| *merrilli* | 163247 | female | 1972 | 03 | 29 | USA | California | Yreka 1 1/2 mi E and 4 mi S, Brazie Ranch, Killgore Hills Rd. | 41.673 | -122.612 |
| *merrilli* | 163249 | male | 1972 | 03 | 29 | USA | California | Yreka 1 1/2 mi E and 4 mi S, Brazie Ranch, Killgore Hills Rd. | 41.673 | -122.612 |
| *merrilli* | 163250 | female | 1972 | 03 | 29 | USA | California | Yreka 1 1/2 mi E and 4 mi S, Brazie Ranch, Killgore Hills Rd. | 41.673 | -122.612 |
| *merrilli* | 163251 | female | 1972 | 03 | 29 | USA | California | Yreka 1 1/2 mi E and 4 mi S, Brazie Ranch, Killgore Hills Rd. | 41.673 | -122.612 |
| *merrilli* | 163254 | female | 1972 | 03 | 29 | USA | California | Yreka 1 1/2 mi E and 4 mi S, Brazie Ranch, Killgore Hills Rd. | 41.673 | -122.612 |
| *merrilli* | 163175 | male | 1971 | 04 | 01 | USA | California | Montague 1 mi NW, Airport Field, Shasta View Ranch | 41.731 | -122.545 |

| | | | | | | | | | | |
|---|---|---|---|---|---|---|---|---|---|---|
| *merrilli* | 163176 | female | 1971 | 04 | 01 | USA | California | Montague 1 mi NW, Airport Field, Shasta View Ranch | 41.731 | -122.545 |
| *merrilli* | 163177 | male | 1971 | 04 | 01 | USA | California | Montague 1 mi NW, Airport Field, Shasta View Ranch | 41.731 | -122.545 |
| *merrilli* | 163179 | female | 1971 | 04 | 02 | USA | California | Montague 1 mi NW, Airport Field, Shasta View Ranch | 41.731 | -122.545 |
| *merrilli* | 163180 | male | 1971 | 04 | 02 | USA | California | Montague 1 mi NW, Airport Field, Shasta View Ranch | 41.731 | -122.545 |
| *merrilli* | 163216 | male | 1972 | 03 | 26 | USA | California | Montague 1 mi NW, Shasta View Ranch | 41.737 | -122.54 |
| *merrilli* | 163217 | female | 1972 | 03 | 26 | USA | California | Montague 1 mi NW, Shasta View Ranch | 41.737 | -122.54 |
| *merrilli* | 163218 | female | 1972 | 03 | 26 | USA | California | Montague 1 mi NW, Shasta View Ranch | 41.737 | -122.54 |
| *merrilli* | 163219 | male | 1972 | 03 | 26 | USA | California | Montague 1 mi NW, Shasta View Ranch | 41.737 | -122.54 |
| *merrilli* | 163220 | female | 1972 | 03 | 26 | USA | California | Montague 1 mi NW, Shasta View Ranch | 41.737 | -122.54 |
| *occidentalis* | 67284 | female | 1934 | 06 | 08 | USA | Arizona | Big Lake, 20 mi S Springerville | 33.889 | -109.413 |
| *occidentalis* | 67286 | female | 1934 | 06 | 08 | USA | Arizona | Big Lake, 20 mi S Springerville | 33.889 | -109.413 |
| *occidentalis* | 87783 | male | 1942 | 06 | 12 | USA | Arizona | U.S. Highway 89, house Rock Ranch, house Rock Valley | 36.718 | -111.955 |
| *occidentalis* | 87784 | male | 1942 | 06 | 12 | USA | Arizona | U.S. Highway 89, house Rock Ranch, house Rock Valley | 36.718 | -111.955 |
| *occidentalis* | 87785 | male | 1942 | 06 | 12 | USA | Arizona | U.S. Highway 89, house Rock Ranch, house Rock Valley | 36.718 | -111.955 |
| *occidentalis* | 87786 | male | 1942 | 06 | 12 | USA | Arizona | U.S. Highway 89, house Rock Ranch, house Rock Valley | 36.718 | -111.955 |
| *occidentalis* | 87788 | female | 1942 | 06 | 12 | USA | Arizona | U.S. Highway 89, house Rock Ranch, house Rock Valley | 36.718 | -111.955 |

| | | | | | | | | | | |
|---|---|---|---|---|---|---|---|---|---|---|
| *occidentalis* | 149571 | female | 1963 | 03 | 12 | Mexico | Chihuahua | Ojo Laguna | 29.462 | -106.33 |
| *occidentalis* | 149572 | male | 1963 | 03 | 28 | Mexico | Chihuahua | Ojo Laguna | 29.462 | -106.33 |
| *occidentalis* | 82390 | male | 1928 | 03 | 08 | USA | New Mexico | Saccatone Cr. | 33.155 | -108.702 |
| *occidentalis* | 71116 | male | 1937 | 03 | 16 | USA | New Mexico | 1 mi S Santa Fe | 35.672 | -105.937 |
| *occidentalis* | 71117 | male | 1937 | 03 | 16 | USA | New Mexico | 1 mi S Santa Fe | 35.672 | -105.937 |
| *occidentalis* | 71118 | male | 1937 | 03 | 16 | USA | New Mexico | 1 mi S Santa Fe | 35.672 | -105.937 |
| *occidentalis* | 71119 | male | 1937 | 03 | 16 | USA | New Mexico | 1 mi S Santa Fe | 35.672 | -105.937 |
| *occidentalis* | 71120 | female | 1937 | 03 | 16 | USA | New Mexico | 1 mi S Santa Fe | 35.672 | -105.937 |
| *occidentalis* | 71121 | female | 1937 | 03 | 16 | USA | New Mexico | 1 mi S Santa Fe | 35.672 | -105.937 |
| *occidentalis* | 82391 | female | 1933 | 05 | 06 | USA | New Mexico | Animas R. | 36.822 | -107.992 |
| *rubea* | 22710 | female | 1912 | 04 | 09 | USA | California | Marysville Buttes, 4 mi NW Sutter | 39.201 | -121.801 |
| *rubea* | 22711 | female | 1912 | 04 | 09 | USA | California | Marysville Buttes, 4 mi NW Sutter | 39.201 | -121.801 |
| *rubea* | 22714 | male | 1912 | 04 | 09 | USA | California | Marysville Buttes, 4 mi NW Sutter | 39.201 | -121.801 |
| *rubea* | 22707 | male | 1912 | 04 | 05 | USA | California | Marysville Buttes, 8 mi NW Sutter | 39.239 | -121.856 |
| *rubea* | 22708 | male | 1912 | 04 | 04 | USA | California | Marysville Buttes, 8 mi NW Sutter | 39.239 | -121.856 |
| *rubea* | 22709 | female | 1912 | 04 | 04 | USA | California | Marysville Buttes, 8 mi NW Sutter | 39.239 | -121.856 |
| *rubea* | 22719 | female | 1912 | 05 | 28 | USA | California | Chambers Ravine, 4 mi N Oroville | 39.57 | -121.555 |
| *rubea* | 22720 | male | 1912 | 05 | 28 | USA | California | Chambers Ravine, 4 mi N Oroville | 39.57 | -121.555 |
| *rubea* | 22721 | female | 1912 | 05 | 28 | USA | California | Chambers Ravine, 4 mi N Oroville | 39.57 | -121.555 |
| *rubea* | 22722 | male | 1912 | 05 | 28 | USA | California | Chambers Ravine, 4 mi N Oroville | 39.57 | -121.555 |
| *rubea* | 22724 | male | 1912 | 05 | 28 | USA | California | Chambers Ravine, 4 mi N Oroville | 39.57 | -121.555 |
| *rubea* | 22727 | female | 1912 | 05 | 28 | USA | California | Chambers Ravine, 4 mi N Oroville | 39.57 | -121.555 |
| *rubea* | 22729 | female | 1912 | 05 | 28 | USA | California | Chambers Ravine, 4 mi N Oroville | 39.57 | -121.555 |
| *rubea* | 64579 | female | 1934 | 03 | 03 | USA | California | Dry Creek, 10 mi NE Oroville | 39.582 | -121.56 |
| *rubea* | 64581 | male | 1934 | 03 | 03 | USA | California | Dry Creek, 10 mi NE Oroville | 39.582 | -121.56 |

| | | | | | | | | | | |
|---|---|---|---|---|---|---|---|---|---|---|
| *rubea* | 64583 | male | 1934 | 03 | 03 | USA | California | Dry Creek, 10 mi NE Oroville | 39.582 | -121.56 |
| *rubea* | 64587 | male | 1934 | 03 | 03 | USA | California | Dry Creek, 10 mi NE Oroville | 39.582 | -121.56 |
| *rubea* | 64590 | female | 1934 | 03 | 03 | USA | California | Dry Creek, 10 mi NE Oroville | 39.582 | -121.56 |
| *rubea* | 64592 | male | 1934 | 03 | 03 | USA | California | Dry Creek, 10 mi NE Oroville | 39.582 | -121.56 |
| *rubea* | 64593 | female | 1934 | 03 | 03 | USA | California | Dry Creek, 10 mi NE Oroville | 39.582 | -121.56 |
| *sierrae* | 165405 | female | 1977 | 07 | 12 | USA | California | Kyburz Flat | 39.502 | -120.236 |
| *sierrae* | 165406 | male | 1977 | 07 | 12 | USA | California | Kyburz Flat | 39.502 | -120.236 |
| *sierrae* | 165408 | female | 1977 | 07 | 12 | USA | California | Kyburz Flat | 39.502 | -120.236 |
| *sierrae* | 165409 | female | 1977 | 07 | 12 | USA | California | Kyburz Flat | 39.502 | -120.236 |
| *sierrae* | 165410 | male | 1977 | 07 | 12 | USA | California | Kyburz Flat | 39.502 | -120.236 |
| *sierrae* | 165411 | male | 1977 | 07 | 12 | USA | California | Kyburz Flat | 39.502 | -120.236 |
| *sierrae* | 165415 | male | 1977 | 07 | 13 | USA | California | Kyburz Flat | 39.502 | -120.236 |
| *sierrae* | 165416 | female | 1977 | 07 | 13 | USA | California | Kyburz Flat | 39.502 | -120.236 |
| *sierrae* | 165417 | female | 1977 | 07 | 13 | USA | California | Kyburz Flat | 39.502 | -120.236 |
| *sierrae* | 165419 | male | 1977 | 07 | 13 | USA | California | Kyburz Flat | 39.502 | -120.236 |
| *sierrae* | 162699 | male | 1971 | 06 | 18 | USA | California | 8 1/2 mi E Mt.Ingalls, SW Corner of McReynolds Valley | 39.97 | -120.46 |
| *sierrae* | 162700 | female | 1971 | 06 | 18 | USA | California | 8 1/2 mi E Mt.Ingalls, SW Corner of McReynolds Valley | 39.97 | -120.46 |
| *sierrae* | 162701 | female | 1971 | 06 | 18 | USA | California | 8 1/2 mi E Mt.Ingalls, SW Corner of McReynolds Valley | 39.97 | -120.46 |
| *sierrae* | 162702 | male | 1971 | 06 | 18 | USA | California | 8 1/2 mi E Mt.Ingalls, SW Corner of McReynolds Valley | 39.97 | -120.46 |
| *sierrae* | 162703 | male | 1971 | 06 | 18 | USA | California | 8 1/2 mi E Mt.Ingalls, SW Corner of McReynolds Valley | 39.97 | -120.46 |

| | | | | | | | | | | |
|---|---|---|---|---|---|---|---|---|---|---|
| *sierrae* | 162704 | female | 1971 | 06 | 18 | USA | California | 8 1/2 mi E Mt.Ingalls, SW Corner of McReynolds Valley | 39.97 | -120.46 |
| *sierrae* | 162708 | male | 1971 | 06 | 18 | USA | California | 8 1/2 mi E Mt.Ingalls, SW Corner of McReynolds Valley | 39.97 | -120.46 |
| *sierrae* | 162711 | female | 1971 | 06 | 18 | USA | California | 8 1/2 mi E Mt.Ingalls, SW Corner of McReynolds Valley | 39.97 | -120.46 |
| *sierrae* | 166913 | male | 1979 | 07 | 10 | USA | California | McReynolds Valley, SW Corner | 39.986 | -120.459 |
| *sierrae* | 166915 | female | 1979 | 07 | 10 | USA | California | McReynolds Valley, SW Corner | 39.986 | -120.459 |
| *strigata* | 46853 | male | 1926 | 03 | 05 | USA | Oregon | 5 mi N Medford | 42.399 | -122.874 |
| *strigata* | 160767 | female | 1971 | 08 | 31 | USA | Oregon | Agate Reservoir, 4 mi ESE White City | 42.413 | -122.772 |
| *strigata* | 160768 | male | 1971 | 08 | 31 | USA | Oregon | Agate Reservoir, 4 mi ESE White City | 42.413 | -122.772 |
| *strigata* | 160769 | female | 1971 | 08 | 31 | USA | Oregon | Agate Reservoir, 4 mi ESE White City | 42.413 | -122.772 |
| *strigata* | 160772 | female | 1971 | 09 | 01 | USA | Oregon | Agate Reservoir, 4 mi ESE White City | 42.413 | -122.772 |
| *strigata* | 160773 | female | 1971 | 09 | 01 | USA | Oregon | Agate Reservoir, 4 mi ESE White City | 42.413 | -122.772 |
| *strigata* | 160777 | female | 1971 | 09 | 01 | USA | Oregon | Agate Reservoir, 4 mi ESE White City | 42.413 | -122.772 |
| *strigata* | 164441 | male | 1976 | 05 | 22 | USA | Oregon | Agate Reservoir, 4 mi ESE White City | 42.413 | -122.772 |
| *strigata* | 164443 | male | 1976 | 05 | 22 | USA | Oregon | Agate Reservoir, 4 mi ESE White City | 42.413 | -122.772 |
| *strigata* | 164445 | female | 1976 | 05 | 22 | USA | Oregon | Agate Reservoir, 4 mi ESE White City | 42.413 | -122.772 |
| *strigata* | 164447 | female | 1976 | 05 | 22 | USA | Oregon | Agate Reservoir, 4 mi ESE White City | 42.413 | -122.772 |
| *strigata* | 164450 | male | 1976 | 05 | 22 | USA | Oregon | Agate Reservoir, 4 mi ESE White City | 42.413 | -122.772 |
| *strigata* | 164452 | female | 1976 | 05 | 22 | USA | Oregon | Agate Reservoir, 4 mi ESE White City | 42.413 | -122.772 |
| *strigata* | 164454 | female | 1976 | 05 | 22 | USA | Oregon | Agate Reservoir, 4 mi ESE White City | 42.413 | -122.772 |
| *strigata* | 164457 | female | 1976 | 05 | 22 | USA | Oregon | Agate Reservoir, 4 mi ESE White City | 42.413 | -122.772 |
| *strigata* | 46851 | male | 1926 | 01 | 27 | USA | Oregon | 3 mi S Eagle Point | 42.429 | -122.802 |
| *strigata* | 65511 | male | 1934 | 05 | 27 | USA | Oregon | 8 mi NNE Medford | 42.434 | -122.815 |

| | | | | | | | | | | |
|---|---|---|---|---|---|---|---|---|---|---|
| *strigata* | 65513 | male | 1934 | 05 | 27 | USA | Oregon | 8 mi NNE Medford | 42.434 | -122.815 |
| *strigata* | 33529 | male | 1896 | 05 | 27 | USA | Oregon | Salem | 44.943 | -123.034 |
| *strigata* | 82381 | male | 1921 | 04 | 24 | USA | Washington | Tacoma | 47.253 | -122.443 |
| *utahensis* | 134099 | male | 1956 | 04 | 12 | USA | Idaho | N of Elba | 42.248 | -113.561 |
| *utahensis* | 67608 | female | 1934 | 06 | 03 | USA | Idaho | 10 mi N Riddle | 42.332 | -116.109 |
| *utahensis* | 67701 | male | 1934 | 06 | 14 | USA | Idaho | 3.5 mi S Decko | 42.468 | -113.627 |
| *utahensis* | 134098 | male | 1956 | 04 | 11 | USA | Idaho | 10 mi N Rupert | 42.764 | -113.676 |
| *utahensis* | 69181 | female | 1936 | 07 | 25 | USA | Idaho | Pahsimeroi Valley, 10 mi S Goldburg | 44.241 | -113.645 |
| *utahensis* | 69184 | female | 1936 | 07 | 25 | USA | Idaho | Pahsimeroi Valley, 10 mi S Goldburg | 44.241 | -113.645 |
| *utahensis* | 69178 | female | 1936 | 07 | 22 | USA | Idaho | Pahsimeroi Valley, 6 mi S Goldburg | 44.299 | -113.645 |
| *utahensis* | 57430 | female | 1930 | 04 | 22 | USA | Nevada | 5 mi SE Millett P.O. | 38.964 | -117.114 |
| *utahensis* | 57431 | male | 1930 | 04 | 22 | USA | Nevada | 5 mi SE Millett P.O. | 38.964 | -117.114 |
| *utahensis* | 57432 | female | 1930 | 04 | 23 | USA | Nevada | 5 mi SE Millett P.O. | 38.964 | -117.114 |
| *utahensis* | 57441 | male | 1930 | 05 | 13 | USA | Nevada | 5 mi SE Millett P.O. | 38.964 | -117.114 |
| *utahensis* | 64678 | female | 1934 | 05 | 19 | USA | Nevada | on Lincoln Hwy., 15 mi SE Eureka | 39.347 | -115.642 |
| *utahensis* | 133192 | female | 1955 | 06 | 11 | USA | Nevada | 9 mi E Jarbidge Pk. | 41.841 | -115.216 |
| *utahensis* | 134250 | male | 1956 | 06 | 05 | USA | Nevada | 4 mi N Jarbidge | 41.932 | -115.431 |
| *utahensis* | 90359 | male | 1942 | 04 | 12 | USA | Utah | 10 mi W Salt Lake Airport | 40.625 | -112.182 |
| *utahensis* | 90360 | male | 1942 | 04 | 12 | USA | Utah | 10 mi W Salt Lake Airport | 40.625 | -112.182 |
| *utahensis* | 90361 | male | 1942 | 04 | 12 | USA | Utah | 10 mi W Salt Lake Airport | 40.625 | -112.182 |
| *utahensis* | 90362 | male | 1942 | 04 | 12 | USA | Utah | 10 mi W Salt Lake Airport | 40.625 | -112.182 |
| *utahensis* | 90365 | female | 1942 | 04 | 18 | USA | Utah | 10 mi W Salt Lake Airport | 40.625 | -112.182 |

Supplementary Table S2: Principal component analysis loadings for plumage data set. The first principal component axis loads positively with brighter, more patterned dorsal plumage, while the second represents a tradeoff between brightness and dorsal plumage.

| | PC1 | PC2 | PC3 |
|---|---|---|---|
| Brightness | 0.53 | 0.85 | 0.00 |
| Achieved Chroma | 0.00 | 0.00 | 1.00 |
| Dorsal Patterning | 0.85 | -0.53 | 0.00 |

Supplementary Table S3: Principal component analysis loadings for soil color data set. The first principal component axis corresponds to soil brightness, while the second principal component axis corresponds to soil redness.

| | PCA 1 Loading | PCA 2 Loading |
|---|---|---|
| Red | 0.70 | 0.64 |
| Green | 0.57 | -0.26 |
| Blue | 0.42 | -0.72 |

Supplementary Table S4: Principal component analysis loadings for soil color data set. The first principal component axis corresponds to soil granularity in which higher scores indicate more coarse particles in the soil.

| Character | PCA 1 Loading |
|---|---|
| Coarse fragments (> 2 mm; volume %) | 0.16 |
| Sand (mass %) | 0.8 |
| Silt (mass %) | -0.49 |
| Clay (mass %) | -0.31 |

Supplementary Table S5: Loadings of principal component analysis of 19 bioclimatic variables used in linear models to examine associations with solar absorption and models of evaporative cooling costs. Bold values contributed strongly (absolute value of loading above 0.1) to the corresponding principal component axis.

| WorldClim Variable | PC1 | PC2 | PC3 |
|---|---|---|---|
| Annual Mean Temperature | 0.02 | 0.06 | **0.33** |
| Mean Diurnal Range | 0.00 | 0.01 | 0.10 |
| Isothermality | 0.00 | 0.01 | 0.01 |
| Temperature Seasonality | **-1.00** | -0.02 | 0.04 |
| Max Temperature of Warmest Month | 0.00 | 0.06 | **0.39** |
| Min Temperature of Coldest Month | 0.03 | 0.05 | **0.31** |
| Temperature Annual Range | -0.03 | 0.01 | 0.08 |
| Mean Temperature of Wettest Quarter | 0.01 | **0.12** | **0.13** |
| Mean Temperature of Driest Quarter | 0.02 | 0.01 | **0.54** |
| Mean Temperature of Warmest Quarter | 0.00 | 0.07 | **0.33** |
| Mean Temperature of Coldest Quarter | 0.03 | 0.06 | **0.32** |

| | | | |
|---|---|---|---|
| Annual Precipitation | 0.01 | **-0.82** | 0.04 |
| Precipitation of Wettest Month | 0.01 | **-0.14** | 0.03 |
| Precipitation of Driest Month | 0.00 | -0.02 | -0.03 |
| Precipitation Seasonality | 0.01 | 0.00 | 0.03 |
| Precipitation of Wettest Quarter | 0.02 | **-0.37** | 0.06 |
| Precipitation of Driest Quarter | -0.01 | -0.06 | -0.08 |
| Precipitation of Warmest Quarter | -0.01 | -0.05 | **-0.20** |
| Precipitation of Coldest Quarter | 0.02 | **-0.36** | **0.21** |

Table S6. Model selection for climatic covariates and reduction in cooling costs.

| **Model** | logLik | AICc | ΔAIC | weight |
|---|---|---|---|---|
| Temperature | 108.416 | -210.5 | 0.00 | 1 |
| Aridity | 79.907 | -153.5 | 57.02 | 0 |
| Seasonality | 69.396 | -132.4 | 78.04 | 0 |
